# The non-typeable *Haemophilus influenzae* major adhesin Hia is a dual function lectin that binds to human-specific respiratory tract sialic acid glycan receptors

**DOI:** 10.1101/2020.05.08.084038

**Authors:** John M Atack, Christopher J Day, Jessica Poole, Kenneth L. Brockman, Jamie R L Timms, Linda E. Winter, Thomas Haselhorst, Lauren O Bakaletz, Stephen J Barenkamp, Michael P Jennings

## Abstract

NTHi is a human-adapted pathogen that colonises the human respiratory tract. Strains of NTHi express multiple adhesins, however there is a unique, mutually exclusive relationship between the major adhesins Hia and HMW1/2. Approximately 25% of NTHi strains express Hia, a phase-variable autotransporter protein, and which has a critical role in colonisation of the host nasopharynx. The remaining 75% of strains express HMW1/2. Previous work has shown that the HMW1 and HMW2 proteins mediate binding to 2,3- and 2,6-linked sialic acid glycans found in the human respiratory tract. Here we show that that the high affinity binding domain of Hia, binding domain 1 (BD1) is responsible for binding to α2,6-sialyllactosamine glycans. BD1 is highly specific for glycans that incorporate the form of sialic acid expressed by humans, *N*-acetylneuraminic acid (Neu5Ac). We further show that Hia has lower affinity binding activity for 2,3-linked sialic acid and that this binding activity is mediated via a distinct domain. Thus, Hia with its dual binding activities functionally mimics the combined activities of the HMW1 and 2 adhesins. In addition, we show that Hia has a role in biofilm formation by strains of NTHi that express the adhesin. Knowledge of the binding affinity of a major NTHi adhesin, and putative vaccine candidate, will direct and inform development of future vaccines and therapeutic strategies for this important pathogen.

**Importance:** Host-adapted bacterial pathogens like NTHi have evolved specific mechanisms to colonize their restricted host niche. Relatively few of the adhesins expressed by NTHi have been characterized as regards their binding affinity at the molecular level. In this work we show that the major NTHi adhesin, Hia, preferentially binds to Neu5Ac-α2,6-sialyllactosamine, the form of sialic acid expressed in humans. The receptors targeted by Hia in the human airway mirror those targeted by influenza A virus and indicates the broad importance of sialic acid glycans as receptors for airway pathogens.

## Introduction

Non-typeable *Haemophilus influenzae* (NTHi) is a human-adapted pathogen, responsible for multiple acute and chronic infections of the respiratory tract, including otitis media (OM) (1) community acquired pneumonia (2), and chronic obstructive pulmonary disease (COPD) exacerbations (3). Each year there are 31 million new cases of the most severe form of OM, chronic suppurative OM, are diagnosed (4), 60% of whom suffer an associated hearing loss. Globally, there are over 700 million cases of acute OM every year (4); in the USA alone, each year there are ∼25 million episodes of acute OM, >13 million antibiotic prescriptions, and public health costs estimated at $3- $5 billion (5, 6). According to WHO estimates, approximately 65 million people have moderate to severe COPD. Over 3 million people died of COPD in 2005, which corresponded to 5% of all deaths globally (7). Invasive disease caused by NTHi has increased significantly in recent years, in part due to vaccines against *Haemophilus influenzae* type b, and *Streptococcus pneumoniae* (8). At present, there is no effective vaccine against NTHi.

NTHi is commonly carried the human nasopharynx asyptomatically. Many bacterial pathogens express outer-surface proteins that target specific host molecules to allow them to adhere to and persist in specific niches in the host. Examples of bacterial adhesins recognising particular host proteins include the type IV pilus of *Neisseria gonorrhoeae*, which recognises host integrins (9); the type IV pilus of NTHi, which recognizes ICAM1 (10); the curli pili of S*almonella enterica*, which binds host TLR2 receptors; and the FimH protein of uropathogenic *Escherichia coli*, which binds to mannosylated glycoproteins (11). Many bacteria also express virulence factors that belong to the auto-transporter protein family. These proteins have a diverse array of functions including adhesion to host surfaces (12). Auto-transporter proteins are characterised by a large barrel-like C-terminal domain which inserts into the outer-membrane, forming a pore through which the N-terminal effector portion passes to reach the extracellular environment (13, 14). NTHi express many autotransporter proteins (15) that fulfil a variety of roles in NTHi pathobiology. One of these autotransporters, Hia, is an adhesin that is expressed by approximately 25% of NTHi strains (16). The remaining ∼75% of NTHi strains express the HMW1/2 proteins (17), which have previously been demonstrated to be involved in adhesion of NTHi to human cells (18). It is unclear why strains encode genes for Hia or HMW but never both. The HMW1 protein binds to host cell glycans as cellular receptors, specifically α2,3-sialyllactosamine (2-3 SLN) (19). We recently demonstrated that HMW2, which is ∼65% identical to HMW1, binds the related glycan α2,6-sialyllactosamine (2-6 SLN), with high specificity for 2-6 SLN containing *N*-acetylneuraminic acid (Neu5Ac), the form of sialic acid expressed by humans (20). Intriguingly, 2-3 SLN is found mainly in the lower human respiratory tract, whereas 2-6 SLN is found throughout the entire respiratory tract, but predominates in the upper airway (21). It has previously been demonstrated that Hia is required for adherence to Chang epithelial cells (22), and we have demonstrated that Hia is required for colonisation of the host nasopharynx (23). However, the cellular receptor for Hia is currently unknown. We hypothesized that Hia may also recognize host-specific glycans found in the human respiratory tract. In the current study we present an investigation to identify and characterized the Hia cellular receptor.

## Results

### Hia is a lectin that recognizes Neu5Ac-α2,6-lactosamine (2-6 SLN-Ac) with high affinity

In order to determine whether Hia had glycan binding activity, we cloned and over-expressed Hia from NTHi strain R2866, in *E. coli* BL21. Heterologous over-expression of Hia in *E. coli* was used previously to investigate Hia binding activity (22). Hia over-expression was confirmed by Western blot and whole cell ELISA (Supplementary Figure 1). The glycan binding ability of *E. coli* strain BL21 cells expressing Hia (BL21-Hia) was compared to wild type BL21 cells, using glycan array analysis. The background binding of BL21 only was subtracted from BL21-Hia in order to deduce the glycans bound in an Hia-dependent manner. A subset of the identified glycans were characterised for their binding affinity to BL21-Hia using surface plasmon resonance (SPR; Table 1). These studies demonstrated that Hia bound to a number of sialylated glycans, with the greatest affinity for Neu5Ac-α2,6-lactosamine (2-6 SLN-Ac), with a disassociation constant (K_D_) of 185 nM. A comparison of the binding affinity of Hia to matched glycan pairs containing either a terminal *N-*acetylneuraminic acid (Neu5Ac; the only form expressed in humans) or *N-*glycolylneuraminic acid (Neu5Gc; which is expressed in most mammals), showed that Hia preferentially binds to structures containing a terminal Neu5Ac (Table 1), with a ∼7-fold preference for 2,6-SLN-Ac over Neu5Gc-α2,6-lactosamine (2,6 SLN-Gc) (185 nM vs 1.39 μM; Table 1). Whilst some binding to 2,3 SLN-Ac (2.03 μM; Table 1) was observed, this occurred with approximately 11-fold lower affinity than for 2-6 SLN-Ac (185 nM).

**Table 1.**
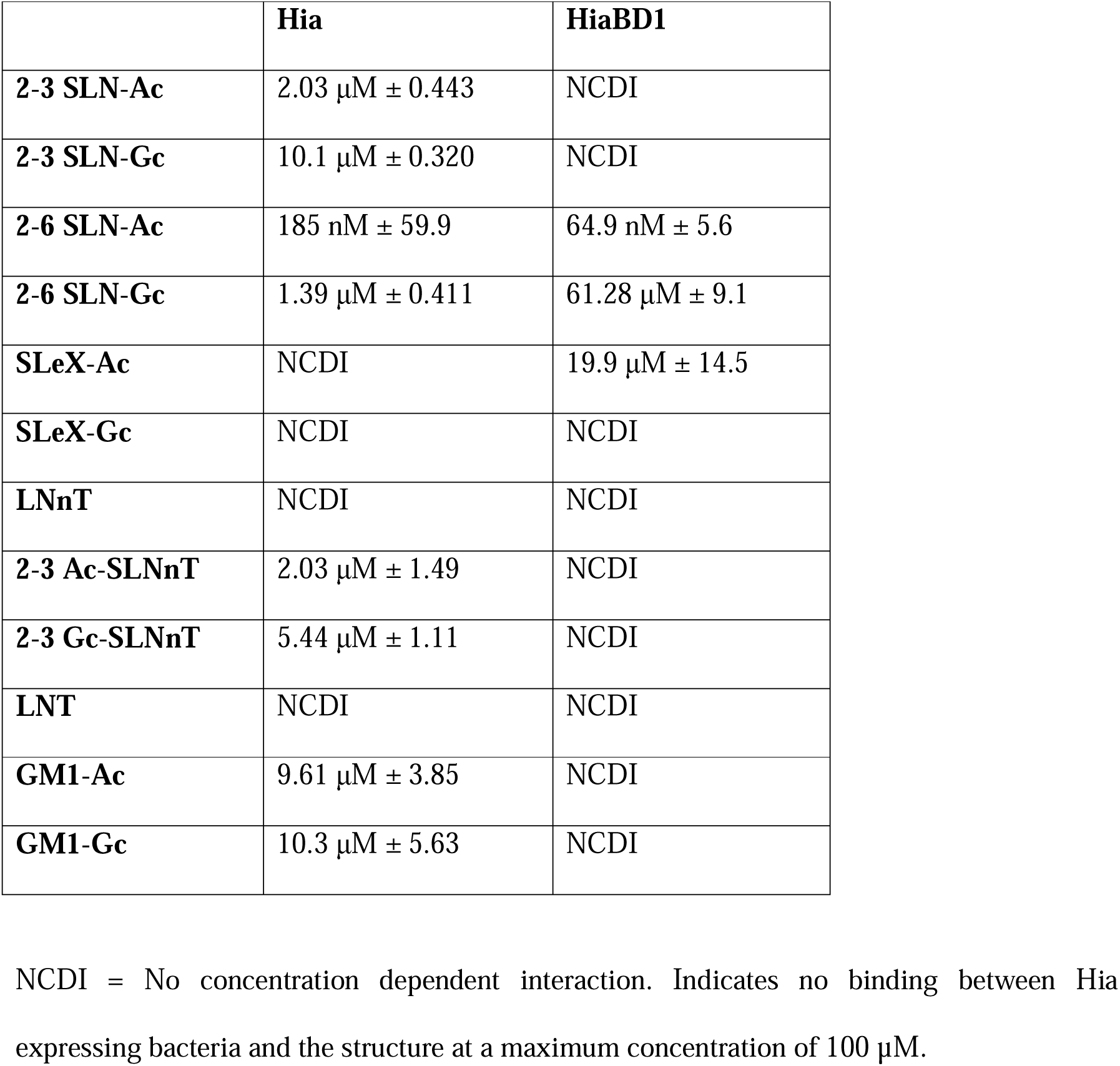
Surface Plasmon Resonance analysis of glycan binding affinity of BL21-Hia and purified recombinant Hia-BD1.

### Modelling shows key interactions between BD1 residues D618 and A620 and the Neu5Ac moiety of 2-6 SLN-Ac

The Hia protein has previously been shown to contain high- and low-affinity host cell binding domains (BD) termed BD1 and BD2 respectively (22, 24). BD1 and BD2 are proposed to bind a common, but unknown cellular receptor (24). Hia BD1 consists of amino acids 541-714 inclusive (22), with residues in the BD1 shown to be essential for binding to Chang epithelial cells when Hia is expressed in *E. coli* (22). To determine the molecular basis of the interactions between Hia BD1 and 2-6 SLN-Ac, we carried out molecular docking studies using the previously published Hia BD1 structure (22). All docking structures of 2-6 SLN-Ac with HiaBD1 indicated interaction of the ligand at the interface of chain A and chain C of HiaBD1. Figure 1 shows a bound structure of 2-6 SLN-Ac that represents a sialic acid-specific binding mode with the negatively charged carboxylate group of the Neu5Ac residue engaging in strong electrostatic interaction with R674. The glycerol side chain of the sialic acid moiety of 2-6 SLN-Ac plays an important role as it engages in hydrogen bonds with D618 and A620. Importantly, the high flexibility of the α(2-6)-linkage of 2-6 SLN-Ac allows the coordination of the lactosamine disaccharide moiety. In addition, our docking studies indicate that residue R674 is involved in coordinating 2-6 SLN-Ac in all potential 25 docked conformations.

**Figure 1.**
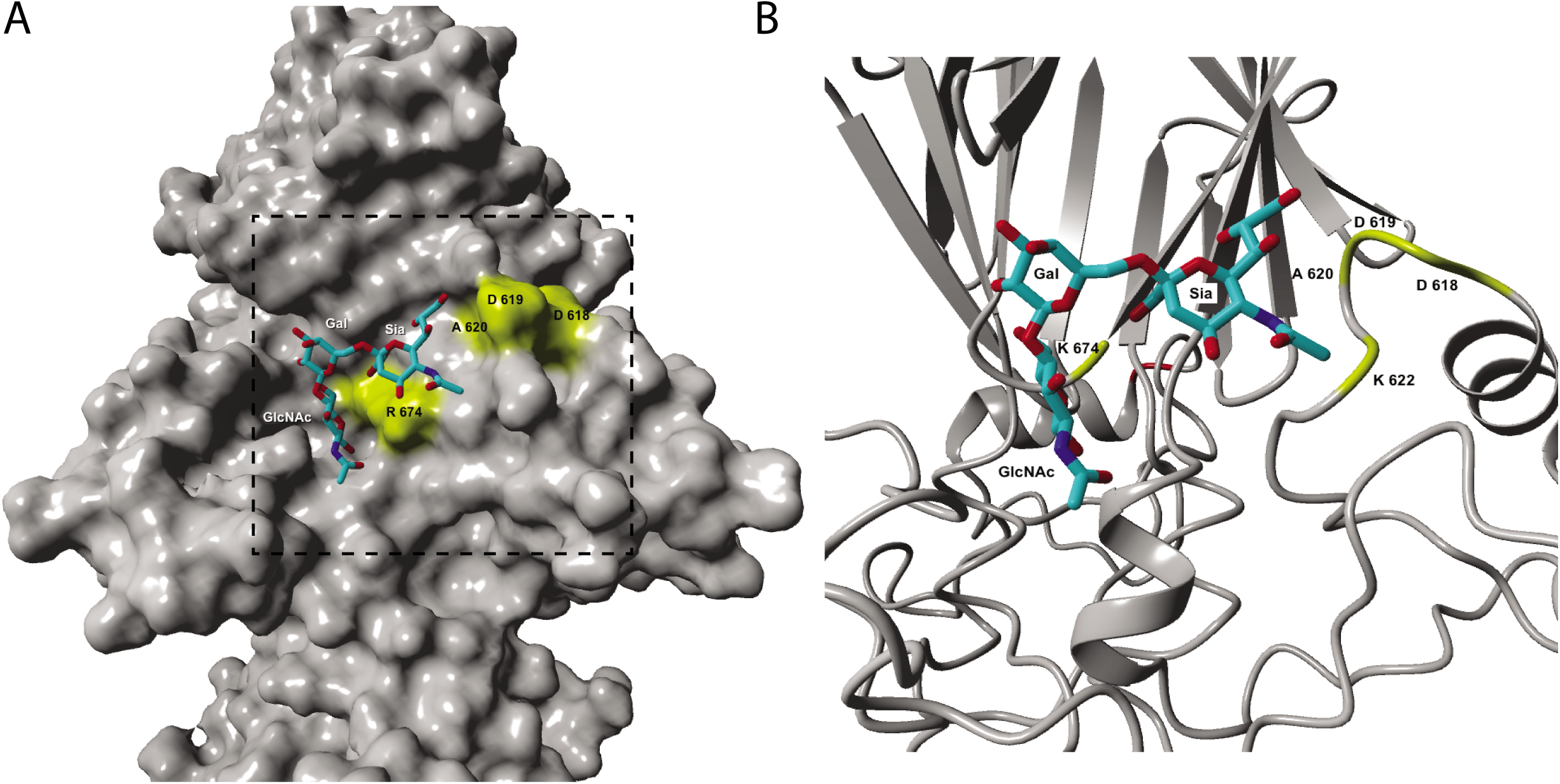
molecular docking of Hia binding domain 1 (BD1) to 2-6 SLN. The previously published structure of Hia BD1 was used (Yeo et al, 2004); PDB accession number 1S7M. Docking structure of 2,6SLN-Ac into Hia (22). **A)** solid surface of 1S7M and **B)** magnified region of 1S7M shown as secondary structure and bound 2-6 SLN-Ac. Key amino acids are labelled.

### Hia BD1 is the site of high affinity interactions with the cellular receptor 2-6 SLN-Ac

Using purified Hia BD1 (aa 514-714 inclusive) (22) we investigated BD1 binding specificity using SPR. Table 1 shows that Hia BD1 binds with high affinity and specificity to 2-6 SLN-Ac, with a K_D_ of 64.9 nM ± 5.6. This value is in a similar range to the affinity we observe with full-length Hia (185 nM ± 59.9). Hia BD1 interacts with 2-6 SLN-Gc with ∼1000 lower affinity (61.28 μM ± 9.1; see Table 1) than with 2-6 SLN-Ac. In order to determine the specific region of BD1 responsible for the interaction with 2-6 SLN-Ac, we constructed a peptide library of BD1 aa 541-714, consisting of peptides of 15 amino acids in length, overlapping consecutive peptides by 10 aa each (for example, peptide one consisted of residues 541-555; peptide two of residues 546-560, etc; Table 2). We used these peptides to block the interaction between BL21-Hia and 2-6 SLN-Ac using an SPR competition assay. Using this methodology, we show that a peptide comprised of twenty amino acid residues containing both D618 and A620 (p16+17; residues 616-635) blocks 100% of the interaction between BL21-Hia and 2-6 SLN-Ac (Table 2). Peptide 16 and Peptide 17 individually result in blocking of 95% and 85% of interactions, respectively, between BL21-Hia and 2-6 SLN-Ac (Table 2). Peptides flanking the region of 16+17 (peptide 15 = aa 611-625; peptide 18 = 626-640; Table 2) only block ∼50% of interactions, with no other peptide 15mer of BD1 blocking interactions between BL21-Hia and 2-6 SLN-Ac (data not shown). Residues D618 and A620 were previously shown to be key for binding to host cells, as when these residues were mutated (D618K and A620R), binding was lost (22). Our blocking studies provide strong evidence that additional residues, and likely secondary structure around these residues that can only form in the 20mer p16+17, mediate direct interaction between 2-6 SLN-Ac and Hia, leading to high-affinity binding.

**Table 2.**
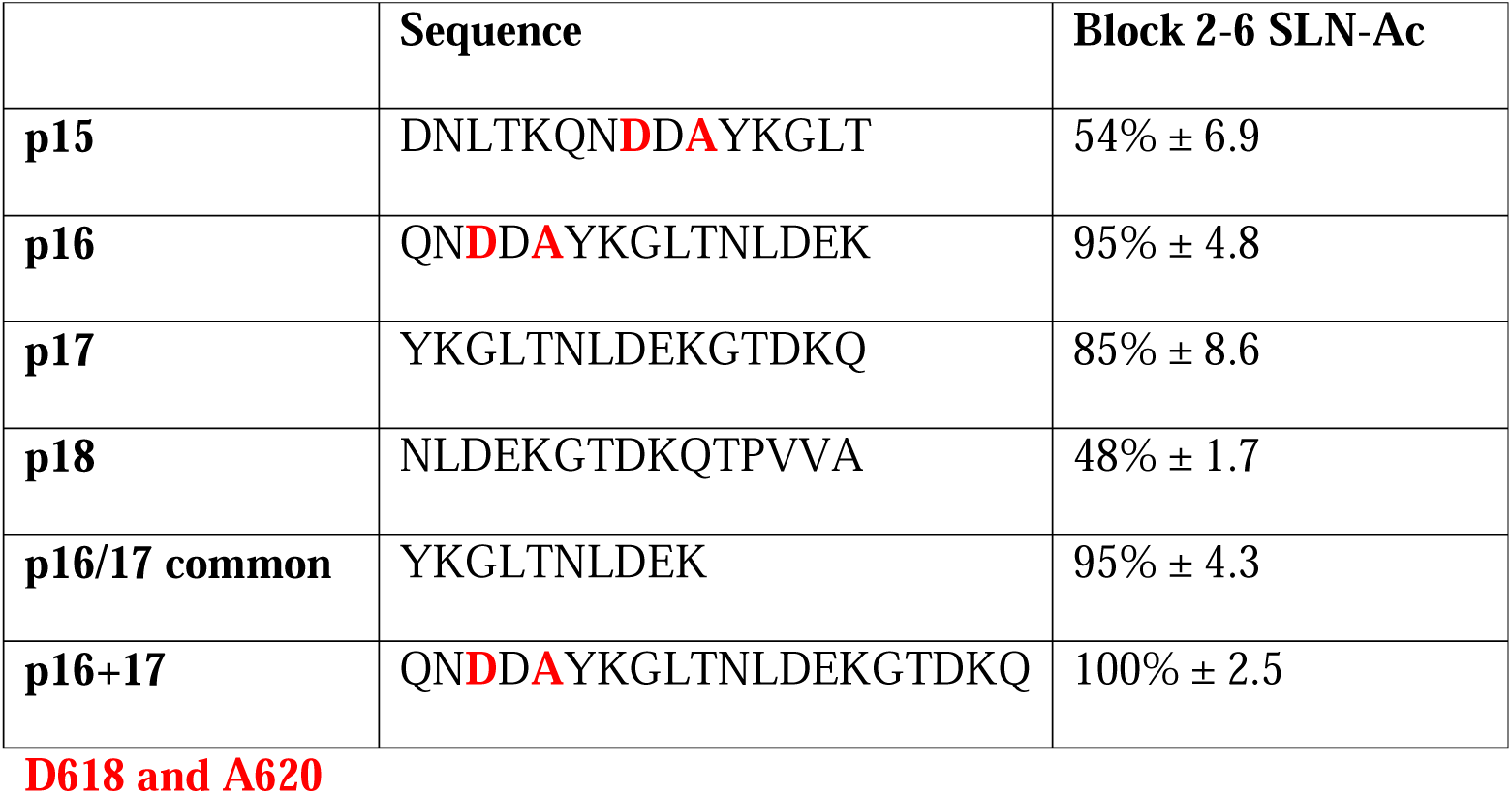
Surface Plasmon Resonance analysis of the blocking activity of peptides to interfer with the Hia : 2-6 SLN-Ac interaction.

In order to confirm these findings, we generated recombinant Hia with the single mutations D618K and A620R, and a double mutant of Hia lacking both of these residues (D618K/A620R double). SPR analysis was used to compare the binding of this panel of Hia mutants with wild type Hia and BD1, using the same subset of glycans (see Table 3). These findings demonstrated that the A620R Hia mutant and the D618K/A620R Hia double mutant (all located in BD1) completely lose the ability to bind 2-6 SLN-Ac, while still maintaining binding to 2-3 SLN-Ac. Collectively, these data demonstrated that the binding site of 2-3 SLN-Ac is not BD1, and confirmed the role of BD1 in binding specificity to 2-6 SLN-Ac.

**Table 3.**
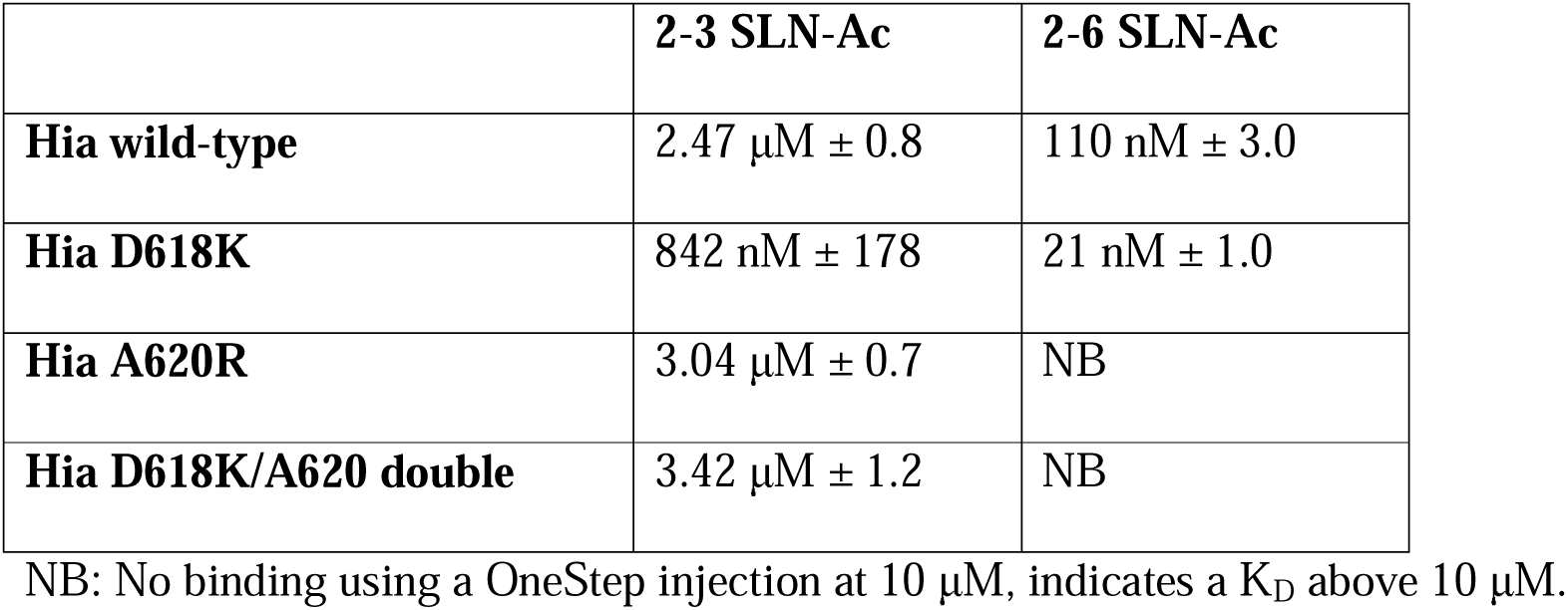
Surface Plasmon Resonance analysis of glycan binding affinity of *E. coli* BL21 expressing wild-type Hia and Hia isogenic mutants.

### Hia is involved in interactions between NTHi and epithelial cells

In order to demonstrate a biological role for Hia in attachment of NTHi to host epithelium, we performed adherence assays using Chang epithelial cells. Prior to carrying out these adherence assays, we confirmed 2-6 SLN was localized on the surface of these cells using Dylight 649 Conjugated SNA, a lectin specific for 2-6 SLN (Figure 2A). Following treatment with sialidase to remove sialylated glycans, 2-6 -SLN was no longer detected on the cell surface by SNA (Figure 2A). Using NTHi strain R2866 that expressed Hia (wild type R2866; R2866 *hia*+), and an isogenic mutant lacking Hia (R2866 *hia::tet*), we showed that the ability of NTHi to adhere to Chang cells decreased when NTHi lacked Hia. Wild type R2866 is unable to bind Chang cells treated with sialidase, which removes sialylated glycan structures (Figure 2B).

**Figure 2.**
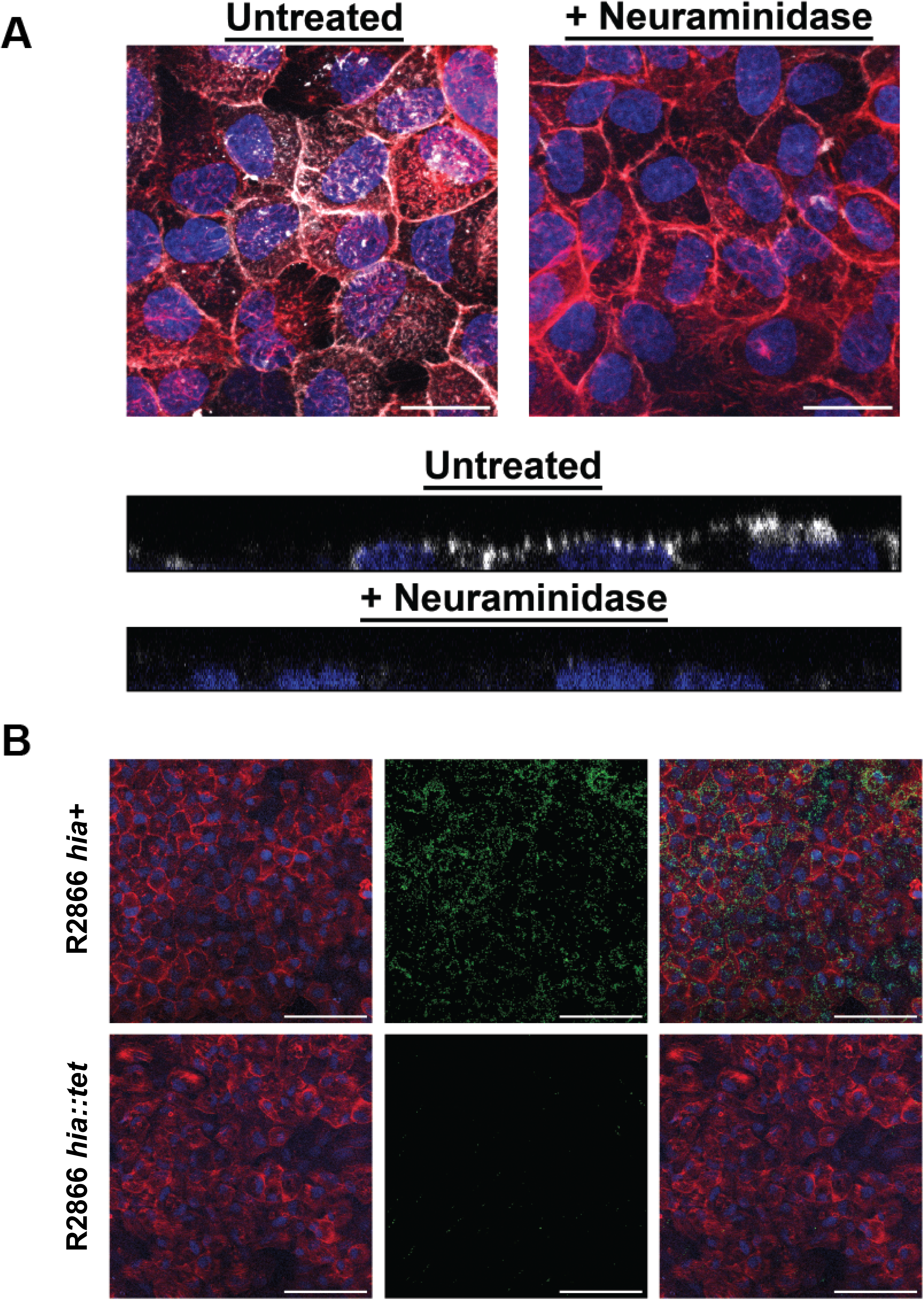
2-6 SLN presence on Chang cells, and adherence of NTHi strain R2866 expressing Hia to Chang cells. **A) 2-6 Sialyl-N-acetyllactosamine expressed on the surface of Chang cells.** Upper panel, top-down view of 2-6 SLN distribution on Change cells or cells pre-treated with neuraminidase. SNAi shown in white, phalloidin shown in red, nuclear DNA shown in blue. Scale bar, 25 µm. Lower panel, representative side view of an optical section through SNAi labelled Chang cells. SNAi shown in white, nuclear DNA shown in blue; **B) Chang cell R2866 adherence.** Adherence of wild type R2866 expressing Hia (*hia*+) and the R2866 *hia::tet* mutant to Chang cells. Left panels, Chang cell monolayer with phalloidin shown in red, nuclear DNA shown in blue. Middle panels, distribution of strain R2866 mutants that constitutively express GFP, shown in green. Bacteria that express Hia (*hia*+) bound markedly better to Chang cells than those that do not express Hia (*hia::tet*). Right panels, merged images that shows distribution of strain R2866 mutants across the surface of the Change cells. Scale bar, 100 µm.

### Residues D618 and A620 are critical to the interaction of Hia with Chang cells

In order to determine the contribution of the key 2-6 SLN-Ac interacting residues (D618, A620), and residue R674, indicated as important from our modelling studies, we carried out adherence assays using a Chang epithelial cell model (23) to determine relative adherence of *E. coli* BL21 strains expressing wild type (wt) Hia and our panel of Hia point mutants. Adherence of *E. coli* BL21 cells to Chang cells was significantly greater when cells expressed wt Hia (14.44% adherence, Figure 3) compared to control cells that did not express Hia (empty BL21; 1.26% adherence; P = 0.0007). BL21 that express the Hia D618K/A620R double mutant exhibited an approximately 4.5-fold decrease in relative adherence (3.21% adherence; P = 0.002) compared to BL21 that express wt Hia. BL21 that expressed Hia R674A, showed an approximate 2-fold decrease in relative adherence compared to cells that express wt Hia (8.4% adherence), but this was not statistically significant compared to cells expressing wt Hia (P = 0.06). These data indicated that the interaction between Hia and 2-6 SLN-Ac is critical to bacterial interactions with epithelial cells, and demonstrate the key contribution of residues D618 and A620 of Hia BD1 in mediating this interaction.

**Figure 3.**
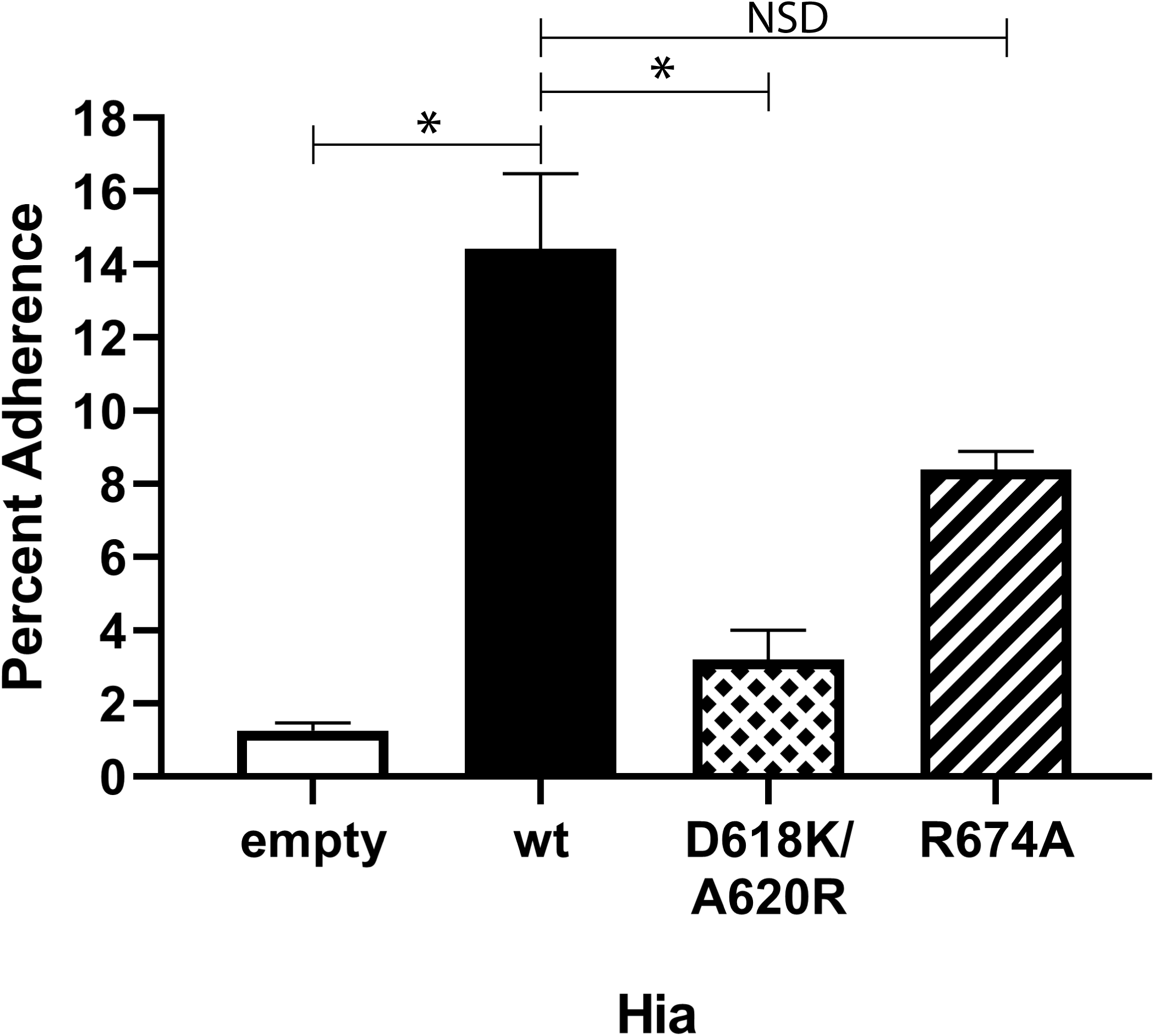
Percent adherence to Chang cells of *E. coli* BL21 strains expressing wild-type Hia or isogenic mutant variants. Percent adherence of each strain is calculated as adherent cfu following 2 hrs incubation / total input cfu. All raw data is presented in Supplementary Data 1. * = P-value = <0.005. NSD = no significant difference. P-values calculated using Student’s t-test

### Expression of Hia in NTHi results in larger more robust biofilms by strains where the hia gene is present

The role of Hia in biofilm formation by two NTHi strains (R2866 and strain 11; both encoding the *hia* gene) was tested using our well defined static biofilm model for NTHi (25). Biofilms were formed for 24 hours at 37°C. Both strain R2866 and strain 11 formed much larger biofilms when Hia was expressed (*hia*+) compared to when it was absent, as assessed by confocal microscopy (Figure 4). NTHi that expressed Hia (*hia*+) formed biofilms with significantly more biomass (*P* = <0.0001 strain R2866 Figure 4A; *P* = <0.01 strain 11; Figure 4B) and were significantly thicker (*P* = <0.0001 strain R2866 Figure 4A; *P* = <0.05 strain 11; Figure 4B) compared to those formed by strains that did not express Hia (*hia::tet*). Descriptively, biofilms formed by NTHi that expressed Hia were significantly denser, and had a lawn-like architecture, compared to those formed by the respective isogenic mutant strain that did not express Hia. Biofilms of the two *hia::tet* isogenic mutant strains were significantly rougher (e.g. had a greater difference in overall surface height and topography) with dense tower-like regions of bacteria surrounded by open water channels (indicated by black in the representative top-down images) (Figure 4). The differences in bacterial distribution within the biofilm are apparent from the area occupied by layer (AOL) graphs (Figure 4). The area occupied by layer is a calculation of the amount (or percentage) of bacterial biomass that is present within each 1-μm optical section of the biofilm taken from the base of the biofilm to the top. These data are plotted wherein the layer closest to the glass surface is at the bottom of the y axis and the top of the biofilm (farthest from the surface) is at the top of the y axis. The relative shift of the blue lines to the right and upward, compared to the red lines (Figure 4), indicated that biofilms formed by NTHi that express Hia are substantially taller and more-dense than those formed by NTHi that do not express Hia. These results indicated that the Hia adhesin was a critical determinant of biofilm structure and organization in these strains, possibly due to increased inter-bacterial associations.

**Figure 4.**
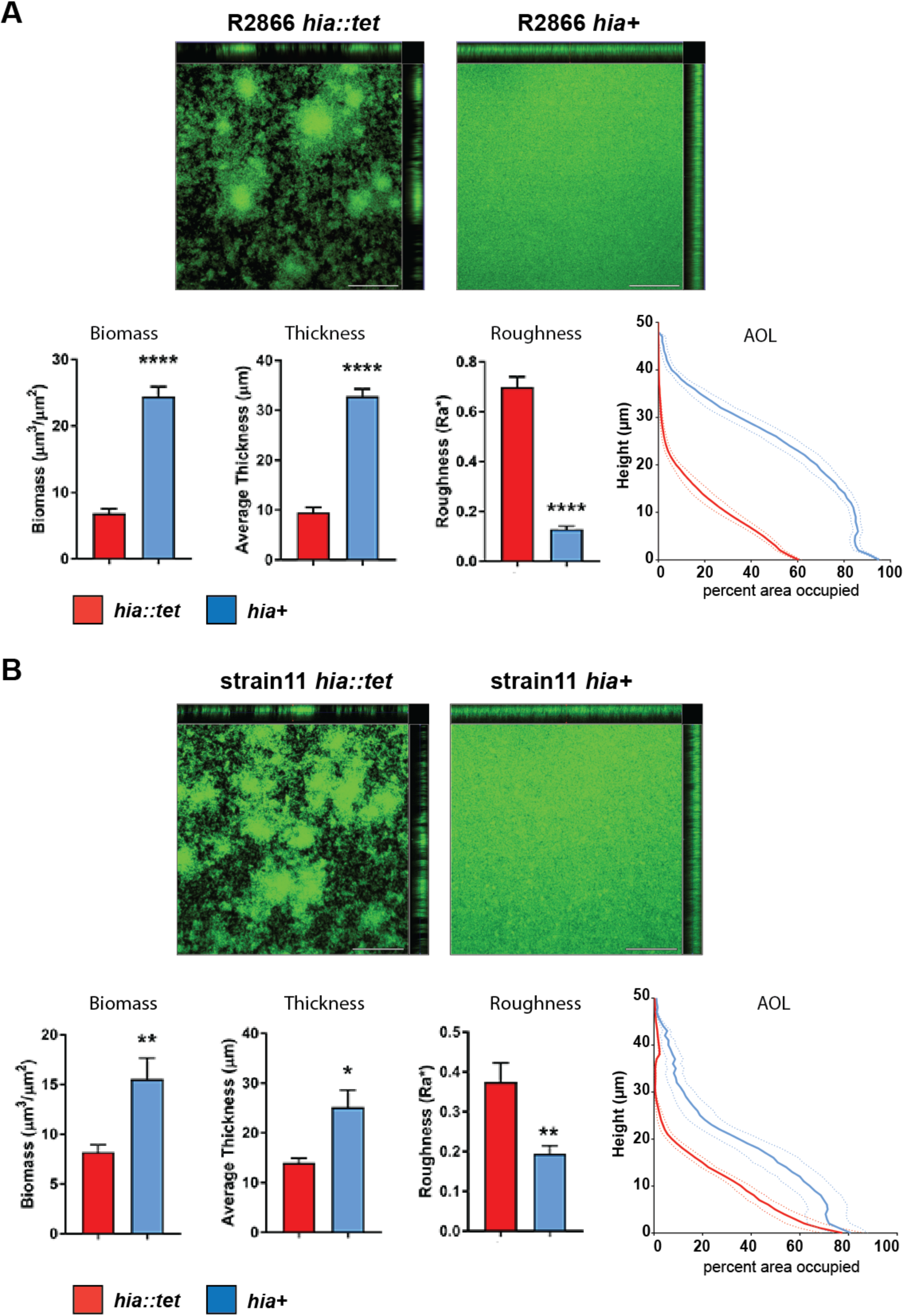
biofilm formation by NTHi strains R2866 and strain 11. Representative orthogonal image renderings of biofilm formation by NTHI strain R2866 and strain 11 *hia*::*tet* and *hia*+ biofilms. Scale bars, 100 μm. Biomass, average thickness and roughness of *hia*::*tet* and *hia*+ biofilms grown for 24 hrs were analyzed by COMSTAT2 and values are shown as mean ± standard error of the mean. *p < 0.05, **p < 0.01, ****p < 0.0001, student’s t-test. Average percent area occupied by bacteria at each individual 1 μm optical section (‘layer’) were determined. Dashed lines indicate standard error of the mean.

## Discussion

In this work, we have demonstrated that the NTHi adhesin Hia is a lectin, with high specificity for host-specific glycans. Hia mediates high-affinity binding to 2-6 SLN-Ac. Molecular modelling studies using the crystal structure of Hia BD1 (22) in complex with 2-6 SLN-Ac showed that Hia residues D618 and A620, and to some extent R674, were critical to this interaction. We experimentally confirmed our modelling using a diverse and comprehensive array of complementary *in vitro* studies. Using a combination of *E. coli* expressing Hia, purified HiaBD1, and a peptide library derived from BD1, we determined that residues D618 and A620 of Hia are required for the high affinity interaction between Hia and 2-6 SLN-Ac. Interestingly, our SPR data also confirmed that Hia recognises 2-3 SLN-Ac, but this interaction was approximately 10-fold lower than that for 2-6 SLN-Ac. However, using the HiaBD1 protein, we confirmed that the interaction of Hia with 2-3 SLN-Ac is not mediated by BD1, which is consistent with previous findings which proposed that Hia contains two binding domains (22). Therefore, contrary to previous work which stated that BD1 and BD2 bind the same ligand (24), we have shown that the two binding domains of Hia interact with distinct ligands; BD1 with 2-6 SLN-Ac and BD2 with 2-3 SLN-Ac. Moreover, in both cases, the preference is for the form of sialic acid expressed by humans (Neu5Ac). This offers an insight into the evolution of NTHi as a human-specific pathogen: although Neu5Gc and Neu5Ac (the precursor to Neu5Gc) can be expressed by most mammals, humans only make Neu5Ac linked glycans, due to a mutation in the CMAH gene responsible for the conversion of Neu5Ac to Neu5Gc (26). As it appears that Hia preferentially binds Neu5Ac linked glycans over Neu5Gc linked glycans, this finding strongly suggests that Hia has evolved to preferentially bind glycans most likely to be present in its human host. Preference for the Neu5Ac form of sialic acid has also been observed in the utilisation of Neu5Ac for macromolecular biosynthesis of bacterial cell surface glycans in NTHi (27), which further supported the central role of sialic acid in the adaptation of NTHi to its human host.

NTHi strains that do not possess the gene encoding *hia* instead encode genes for and express the adhesins HMW1 and HMW2 (18). Previous work has demonstrated that NTHi strains either encode genes for Hia or HMW1/2, but never both, with approximately 75% strains expressing HMW1/HMW2, and the remaining 25% expressing Hia. We have recently demonstrated that HMW2 preferentially binds 2-6 SLN-Ac (20), whereas HMW1 has a preference for 2-3 SLN structures (19). However, HMW1 showed no preference for either Neu5Ac or Neu5Gc containing structures, and had a much lower affinity than HMW2 for 2-6 SLN-Ac (20). Therefore two distinct NTHi adhesins, Hia and HMW1/2, that show a discrete lineage distribution in the NTHi population, have evolved to bind the same subset of glycans: Hia binds 2-6 SLN-Ac preferentially over 2-3 SLN-Ac; HMW2 specifically binds 2-6 SLN-Ac; HMW1 binds a broader range of 2-3 and 2-6 linked glycan structures compared to HMW2, but with lower overall affinity (Figure 5). Production of both 2-6 SLN-Ac and 2-3 SLN-Ac linked glycan structures is found throughout the human airway, but these receptors are not evenly distributed throughout the upper and lower airway; 2-6 SLN-Ac is found in both the upper and lower respiratory tract, while 2’-3’ SLN-Ac linked glycans are found predominantly in the lower respiratory tract (28-30). Thus, NTHi strains that express Hia or HMW1/2 are able to adhere to the entire human respiratory tract. It would be interesting to study the distribution of NTHi strains expressing only HMW1 or HMW2 – based on their differing binding affinities, it may be that HMW2-only expressing strains are more prevalent in upper respiratory tract infections, whereas HMW1-only expressing strains are more disposed to infecting the lower respiratory tract. It is also intriguing to note that the binding specificity of Hia (and HMW1/2) mirrors perfectly that of human influenza A viruses (31-33), indicating that specificity for human specific glycans has evolved in both viral and bacterial human-adapted airway pathogens. A common strategy to block these interactions may therefore serve as a general therapy for both these types of infection. It is well known that infection with the influenza virus predisposes individuals to colonization and infection by *Streptococcus pneumoniae*; although there are likely multiple aspects behind the increased severity of pneumococcal disease following influenza virus infection (34), it is thought that desialylation of the host epithelia by the viral neuraminidase allows for more efficient colonization by the pneumococcus (35). This desialylation in turn increases the susceptibility of these patients to pneumococcal pneumonia following influenza virus infection. The lethality of the 1918 ‘Spanish flu’ outbreak was mainly due to secondary infections by bacterial pathogens, including both *S. pneumoniae* and *H. influenzae*, with up to 95% of the deaths from this pandemic attributable to secondary bacterial infections (36, 37). Thus, the sharing of common receptors indicates the possibility of direct interaction between influenza virus and NTHi occurs during co-infection, and may provide a fruitful area of investigation in the study of the dynamics of this polymicrobial interaction.

**Figure 5.**
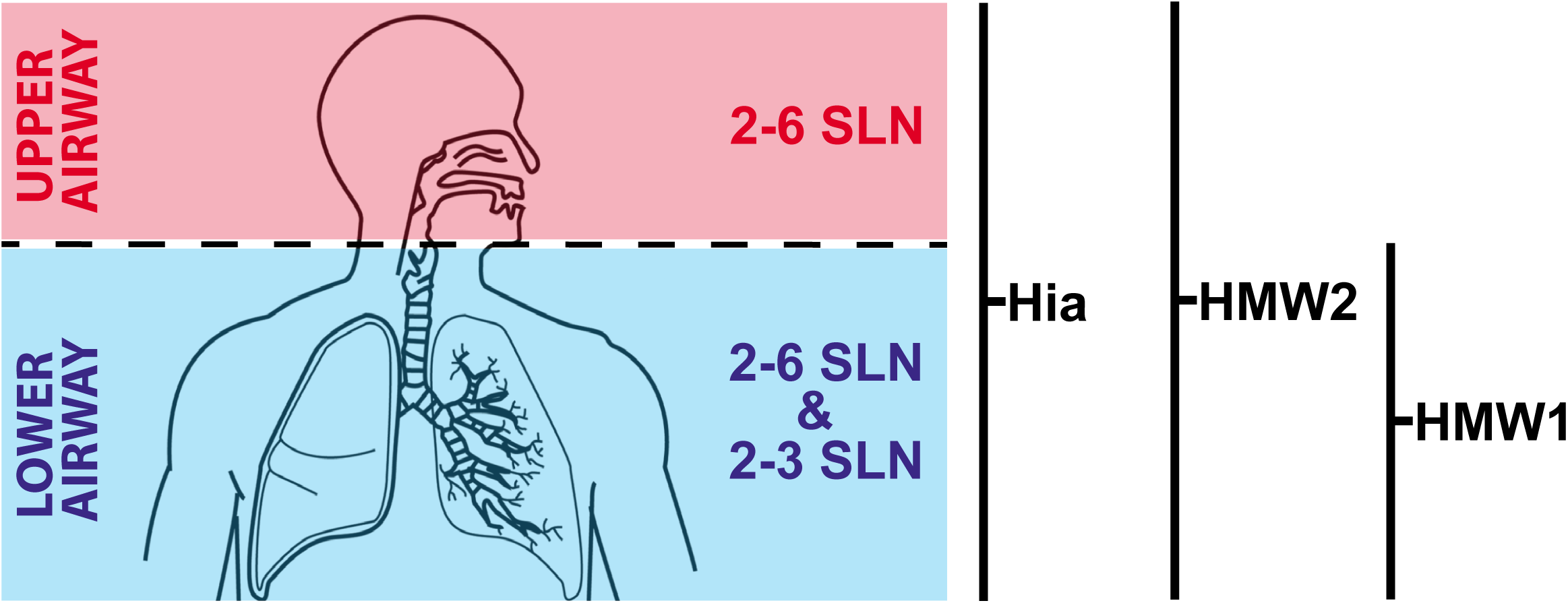
illustration showing distribution of 2-6 SLN and 2-3 SLN in human airway. Hia and HMW2 both bind 2-6 SLN with high affinity, with both showing a very high preference for the human specific sialic acid Neu5Ac (2-6 SLN-Ac) over Neu5Gc (2-6 SLN-Gc). This means NTHi strains expressing either Hia or HMW2 are able to colonise the entire respiratory tract (upper in red, lower in blue). HMW1 preferentially binds 2-3 SLN, and with no affinity for Neu5Ac over Neu5Gc, which means NTHi strains only expressing HMW1 may have a preference for the lower respiratory tract (blue). Schematic diagram taken from GetDrawings.com (http://getdrawings.com/respiratory-system-with-label-drawing#respiratory-system-with-label-drawing-13.jpg) under a CC BY-NC 4.0 Licence.

As well as playing a key role in host colonisation through recognition of human specific glycan structures, we have demonstrated that Hia also plays a key role in biofilm formation by NTHi. Biofilm formation by NTHi has been shown to increase the resistance of bacteria to antibiotics (38), and killing by neutrophils (39) when compared to planktonic counterparts. Increased resistance of bacteria within biofilms to antibiotics has been demonstrated for a number of major human pathogens, such as *Pseudomonas aeruginosa* (40) and *Staphylococcus aureus* (41). Biofilm formation also plays a key role in NTHi disease pathologies, such as middle ear infections (42) and exacerbations of cystic fibrosis (43). Using two diverse strains of NTHi, we showed that Hia is a critical determinant of biofilm development and structure and that the potential to block Hia function through knowledge of its specific binding affinities could play a key role in targeting biofilm formation during disease caused by NTHi.

To summarize, we have provided an in-depth characterization of the binding affinity of the NTHi adhesin Hia, by determining the major human cellular receptors it has evolved to bind and by demonstrating the molecular basis of these interactions. We also demonstrate that Hia has a role in biofilm formation by NTHi, and therefore likely contributes to antibiotic resistance and chronicity by this mechanism. Knowledge of the factors required by NTHi to colonise and cause disease will be key to developing both vaccines and treatments against this organism. Our demonstration that the major NTHi adhesins HMW1 and HMW2 bind the same host glycans as Hia (20), and that these adhesins are expressed by nearly 100% of NTHi strains is a key step towards the development of a rationally designed vaccine against NTHi, and to the production of novel treatments against this pathogen.

## Materials and methods

### Bacterial strains and growth conditions

NTHi strains expressing Hia have been described previously [R2866 (44) and strain 11 (45)]. NTHi strains were routinely grown in Brain-Heart Infusion (BHI) broth supplemented with 1% hemin and 20 µg NAD^+^/mL (sBHI), and grown aerobically at 37°C with 150 rpm shaking. For solid media, 1.5% agar was added to sBHI broth. sBHI media were supplemented with tetracycline (5 µg/mL) as required. Plates were grown at 37°C in atmosphere containing 5% CO_2_. *Escherichia coli* were grown using Luria-Bertani (LB) media at 37°C, and supplemented with tetracycline (5 µg/mL) as required.

### Generation of a hia knockout mutant in NTHi strains R2866 and 11

A region of NTHi R2866 chromosome containing the *hia* promoter and the ATG start and 5′□ region of the gene were generated by PCR using primer pair hia-UP-F / hia-UP-R, and cloned into pGEM Teasy according to manufacturer’s instructions (Promega) to generate plasmid vector Teasy::hiaUP. Inverse PCR was used to linearise this vector at the *hia* start codon using primers hia-INV-F / hia-INV-R. A tetracycline resistance cassette, encoding *tetM*, was generated from plasmid vector pGEM-TetM(B) using M13F and M13R primers.

This was cloned into the linearised Teasy::hiaUP vector so the gene was in the same orientation as the *hia* gene, and orientation confirmed using PCR and sequencing. This vector was designated Teasy::hiaUP::TetM. Following linearization with NgoMIV (New England Biolabs), DNA was transformed into NTHi strains R2866 and strain 11 using the MIV method (46). Transformants were selected on BHI media containing 5 µg tetracycline /mL, and positive colonies confirmed by sequencing and Western blotting using an anti-Hia monoclonal antibody 1F4 (47). Strains were designated as R2866 or strain 11 *hia::tet.*

### Cloning and over-expression of full length Hia in E. coli

Primers HiaFULL-F and HiaFULL-R (Supplementary Table 1) were used to amplify full length wild-type *hia* (R2866_0725) including the signal sequence (residues 1-49) from genomic DNA prepared from NTHi strain R2866. PCR was carried out using KOD hot-start polymerase (EMD Millipore) according to manufacturer’s instructions. Following digestion with BspHI and XhoI (NEB) and clean up, DNA was cloned into pET15b digested with NcoI and XhoI. the resulting plasmid was designated pET15b::Hia. Following confirmation of correct clones by sequencing, over-expression was carried out in *E. coli* BL21 following by inducing cells with 0.5 mM IPTG overnight at 37°C with 200 rpm shaking. Over-expression was confirmed by Western blot as previously described (23) using anti-Hia monoclonal antibody 1F4 (47). Whole cell ELISA using standard methods (48) with modifications as previously described (23) and starting with 1:10,000 dilution of primary antibody anti-Hia monoclonal antibody 1F4 confirmed the location of Hia at the cell surface.

### Generation of Hia point mutants for over-expression

Inverse PCR was carried out using primer pairs designed to introduce point mutations as previously described and used here to abrogate binding of *E. coli* expressing Hia to Chang cells (22). D618K, A620A and a 618/620 double mutant were generated using specific forward primers Hia-D618K-F, Hia-A620R-F, or Hia-618/620-double-F and common reverse primer Hia-618/620-R. A R674A mutant was generated using primer pair Hia-R674A-F and Hia-R674A-R. All inverse PCR reactions were carried out using KOD hot-start polymerase (EMD Millipore) according to manufacturer’s instructions, and a plasmid mini-prep (Qiagen) of pET15b::Hia as template. All primer sequences are listed in Supplementary Table 1. Clones were sequenced using primers either side of the point mutation Hia-screen-F and Hia-screen-R using BigDye 3.1 according to manufacturer’s instructions (Thermo Fisher), and sequenced at Australian Genome Analysis Facility (AGRF, Brisbane, Australia). Over-expression was carried out as described above for Hia wild-type, and cell surface localization confirmed using whole cell ELISA as above.

### Over-expression and purification of Hia BD1

Primers to clone Hia binding domain 1 (BD1; amino acid residues 540-714) were designed based on those from (22). HiaBD1-F and HiaBD1-R were used to amplify BD1 from NTHi strain R2866 genomic DNA using KOD hot-start polymerase (EMD Millipore) according to manufacturer’s instructions. Following digestion with NdeI and BamHI (NEB) and clean up, DNA was cloned into pET15b digested with the same enzymes. This strategy would clone the gene in frame with an N-terminal 6xHis tag for purification. The resulting plasmid was designated pET15b::HiaBD1. Over-expression was carried out in *E. coli* BL21 following by inducing cells with 0.5 mM IPTG overnight at 37C with 200 rpm shaking. Cells were pelleted, resuspended in 1× binding buffer (50 mM NaPO4, 300 mM NaCl, pH7.4), lysed using 0.1 mm glass beads and a Tissue lyser (Qiagen) for 30 mins at 50 osc^-1^ min^-1^. Purification was carried out using TALON gravity flow resin in 1× binding buffer. Protein was eluted from the resin using step wise concentrations of imidazole in 1x binding buffer (10-500mM imidazole), fractions analysed by SDS PAGE, and fractions containing pure BD1 pooled and concentrated using centrifugal concentrators (Millipore, 10kDa cut-off). Pure concentrated BD1 was buffer exchanged into 1x phosphate buffered saline (1x PBS) using the same centrifugal concentrators. Protein was analysed by SDS PAGE, and quantified using an extinction coefficient of 8480 M^-1^ cm^-1^ and MW of 20471.43 Da (based on the sequence of Hia BD1+6xHis tag)

### Glycan array

Glycan array slides were printed using OPEpoxy (CapitalBio) activated substrates with the glycan library as previously described (49) using an ArrayIt Spotbot Extreme 3 contact printer with solid metal pins. The glycan array binding experiments were performed and analysed as previously described (50). Briefly, 1 mL of OD_600_ 0.2 *E. coli* BL21 with heterologous expression of Hia wild-type or Hia point-mutants in PBS were incubated with 15μL of 50 μM Bodipy methylester for 15 minutes, centrifuged at 900g for 3 minutes and the pellet washed 3 times with PBS to removed excess dye. The cell pellet was resuspended in 1 mL of array PBS (1x PBS containing 1 mM CaCl_2_ and 1 mM MgCl_2_) and 300 μL was applied to the slide in a 65 µL gene frame without a coverslip. Slides were washed three times for 2 minutes in array PBS, dried by centrifugation and scanned and analysed using the Scan Array Express software package (Perkin Elmer) and Microsoft Excel for statistical analysis. Binding of Hia was defined as both above the background of the slide (cut off of 550 fluorescence units) and 2-fold and significantly (*p*<0.05) above the background of empty vector BL21 binding to the array by Student’s unpaired t-test of fluorescence of BL21 empty vector controls versus Hia expressing strains of BL21. All glycan array binding data is presented in Supplementary Table 2 and the MIRAGE compliant information is listed in Supplementary Table 3.

### Surface Plasmon Resonance (SPR)

Surface plasmon resonance (SPR) experiments of the full length wild-type Hia expressed on the surface of *E. coli* BL21was performed using a GE Biacore T100 system and a Series S C1 sensor chip using a modification of methods previously described (51, 52). *E. coli* (BL21 strains expressing full length wild-type Hia, point mutants, or BL21 only) cells at 1×10^6^ bacteria/mL were immobilised to the chip surface following the C1 NHS/EDC method template with a contact time of 900 seconds at a flow rate of 5 µL/minute in 10 mM sodium acetate pH 5.5. Interaction of glycans with the bacteria was performed using five-fold serial dilutions with maximum concentration of 20 μM on first analysis and 5 μM when affinities were better defined using single cycle kinetics in 1x PBS pH 7.4 at 20 µL/minute with 60 second contact time and a final dissociation time of 10 minutes. A blank ethanolamine immobilisation was used as a control flow cell and 1x PBS pH 7.4 was used as the zero-concentration control. Regeneration of the bacterial surface was performed by flushing 10 mM Tris 1mM EDTA over the surface for 5 minutes at 30 µL/minute. Affinities (K_D_) were determined using the Biacore T100 evaluation software analysis of double baseline subtracted data. All interactions were measured in triplicate and displayed plus/minus 1 standard deviation of the measured mean.

Purified Hia BD1 protein was immobilised onto a CM5 sensor chip amine capture on a Biacore T100 with a contact time of 600 seconds at a flow rate of 5 µL/minute in 10 mM sodium acetate pH 4.5. Glycans were run at the optimised concentrations outlined above. With the analysis performed as outlined above.

Peptide binding region identification was performed using a modified version of a previously described method (53), competition assays using immobilised Hia expressing cells and flowed peptides and glycan. *E. coli* BL21 expressing full-length wild-type Hia were immobilised onto a H1 sensor chip using a ForteBio Pioneer using a contact time of 720 seconds at a flow rate of 10 µL/minute in 1x PBS at 1×10^8^ bacteria/mL Assays were set up using the NextStep injection feature as previously described (54, 55) with combinations of 2,6-SLN, Hia overlapping peptides and PBS as the negative control used to determine the Hia region that interacted with 2,6-SLN. Analysis was performed using the QDat analysis software package.

### Distribution of 2,6-SLN on Chang cells

Chang cells (1×10^4^ cells) in 100 µL total volume were seeded into Transwell inserts with a 6.5 mm diameter and 0.4 um pore size (Corning Incorporated, Corning, New York). Cell culture medium (DMEM, 10% heat-inactivated calf serum, 2 mM L-glutamine) was replaced daily until cells reached confluence, 2 to 3 days. The apical surface of the cells was rinsed twice with sterile DPBS and then incubated with 0.1 units of neuraminidase (Sigma) in 100 µL DPBS or with DPBS alone for 2 hours at 37°C. The cells were then rinsed twice with DPBS and 100 µL of 10 µg/mL Dylight 649 Conjugated *Sambucus nigra* (EY Laboratories, San Mateo, California) was added to the apical surface and incubated for 15 minutes. The cells were rinsed twice with DPBS, incubated with 3 units of Alexa Fluor™ 594 Phalloidin (Thermo Fisher Scientific, Waltham, Massachusetts) for 30 minutes and rinsed twice. The membrane of the Transwell was excised and then mounted with ProLong™ Glass Antifade Mountant with NucBlue™ Stain (Thermo Fisher Scientific, Waltham, Massachusetts). Images were captured on a LSM 700 laser scanning microscope and rendered with Zeiss Zen software (Zeiss).

### Biofilm formation

Biofilms were formed by NTHI cultured within chambers of eight-well-chambered coverglass slides (Thermo Scientific, Waltham, MA) as described previously (56). Briefly, mid-log phase cultures of NTHI strains were diluted with sBHI. NTHI were inoculated at 4 × 10^4^□c.f.u. in 200□μl total volume per well and slides were incubated at 37°C with 5% atmospheric CO2. Biofilms were grown for a total of 24 h, with the growth medium replaced after 16 h. To visualize, biofilms were stained with LIVE/DEAD BacLight stain (Life Technologies) and fixed overnight in fixative (1.6% paraformaldehyde, 2.5% glutaraldehyde, 4% acetic acid in 0.1□M phosphate buffer, pH 7.4). Fixative was replaced with saline before imaging with a Zeiss 510 Meta-laser scanning confocal microscope. Images were rendered with Zeiss Zen software.

### Analysis of biofilm formation and architecture

Z-stack images acquired at 63x with a Zeiss 510 Meta-laser scanning confocal microscope were analyzed by COMSTAT2 to determine biomass (μm3/μm2), average thickness (μm), roughness (Ra) and percent area occupied by layers. Area occupied by layer was plotted as percent bacterial biomass coverage per 1 μm optical section from the base of the biofilm. Standard error of the mean for replicate biofilms was calculated for each individual layer with GraphPad Prism version 6.0 (GraphPad Software, San Diego, CA).

### Adherence of NTHI strain R2866 to Chang cells

Chang cells (1×10^4^ cells) in 100 µL total volume were seeded into Transwell inserts with a 6.5 mm diameter and 0.4 um pore size (Corning Incorporated, Corning, New York). Cell culture medium (DMEM, 10% heat-inactivated calf serum, 2 mM L-glutamine) was replaced daily until cells reached confluence, 2 to 3 days. The apical surface of the cells was rinsed twice with sterile DPBS. Strain R2866 strains were added to the apical surface of the Chang cells at an MOI of 100 in 50 µL of DPBS and incubated for 30 minutes at 37°C. The cells were rinsed twice with DPBS, incubated with 3 units of Alexa Fluor™ 594 Phalloidin (Thermo Fisher Scientific, Waltham, Massachusetts) for 30 minutes and rinsed twice. The membrane of the Transwell was then excised and mounted with ProLong™ Glass Antifade Mountant with NucBlue™ Stain (Thermo Fisher Scientific, Waltham, Massachusetts). Images were captured on a LSM 700 laser scanning microscope and rendered with Zeiss Zen software (Zeiss).

### Adherence assays with BL21 strains

*E. coli* BL21 strains expressing either wild-type Hia, D618K/A620R double mutant, R674A mutant, or containing the empty pET15b expression vector were grown to mid-log (OD_600_ = ∼0.6) in LB broth containing ampicillin (100 μg/mL), and cfu calculated by serially diluting. Approximately 5×10^5^ cfu (100 μl) each mid-log culture were added to wells of a 24 well plate containing differentiated Chang cells as described previously (23). Six wells were used per strain per experiment. Following addition of BL21, plates were incubated at 37°C for 2 hours. Following incubation, supernatant was removed, and non-adherent cells removed by gentle washing with 1x PBS four times. Adherent cells were released by incubating with 0.05% Trypisn in 1x PBS for 10mins at room-temperature. Adherent bacteria were quantified by serial dilution and plating. Percent adherence was calculated as the number of adherent cfu in relation to the total input cfu per strain. Student’s t-test was carried out using total number of each output and comparing to wild-type cells. All data (total cfu and percent adherence) is presented as Supplementary Data 1.

### Modelling interaction of HiaBD1 with 2-6 SLN

Docking of 2-6 SLN to HiaBD1 was performed using the AutoDock Vina protocol (57) that has the highest scoring power among commercial and academic molecular docking programs (58) and is implemented in the YASARA Structure molecular modelling package (Ver. 16.46) (59). The docking experiment was set up by using the X-ray crystal structure of HiaBD1 (pdb code 1S7M (22) with a box centered D620 using a grid size of 50 Å ⍰ 50 Å 50 Å (x, y, z) covering chain C. A total number of 25 Vina docking runs were performed. The 3D topology of the 2-6 SLN glycan was generated using the carbohydrate builder available at GLYCAM-Web server (http://glycam.org) (60).

## Acknowledgements

Work was funded by the Australian National Health and Medical Research Council (NHMRC) Program Grant 1071659 and Principal Research Fellowship 1138466 to MPJ, Project Grant 1099279 to JMA, an Australian Research Council Discovery Project 180100976 to JMA, a Garnett Passe and Rodney Williams Grant-in-Aid (Supplementation) to JMA, and a National Institutes of Health (NIH; USA) R01 Grant DC015688 to LOB and MPJ. We thank Patrick Azzari for a critical readthrough of, and valuable input to, the manuscript.

---

**Supplementary Figure 1 A)** Western blot showing the over-expression of wild-type Hia in *E. coli* BL21; and **B)** Whole cell ELISA showing that Hia expressed in *E. coli* BL21 is located on the bacterial cell surface

