## Supplementary Table 1 for "The non-typeable *Haemophilus influenzae* major adhesin Hia is a dual function lectin that binds to human-specific respiratory tract sialic acid glycan receptors"

**Supplementary Table 1 - Primers used in this study**

| **Primer** | **Sequence** |
| --- | --- |
| Hia-UP-F | GTAGAAAACTTAGCAACATTAAACGG |
| Hia-UP-R | CCA TTT TGA CCA TTA GCA TCG G |
| Hia-INV-F | GTTATTTGGAATGTTGTGACTCAAA |
| Hia-INV-R | GAA AAA CAA ACA TTT ACA CAA AAA TCA AAT ATT TTC |
| HiaFULL-F | AGTCAG TCATGA ACAAAATTTTTAACGTTATTTGGAATG |
| HiaFULL-R | AGTCAG CTCGAG TTACCACTGGTAACCAACACC |
| Hia-D618K-F | GACAA C TTA ACG AAA CAA AAT **AAA** GAT GCC TAT AAA GGC TTG ACC AAT TTG G |
| Hia-A620R-F | GACAA C TTA ACG AAA CAA AAT GAC GAT **CGC** TAT AAA GGC TTG ACC AAT TTG G |
| Hia-618/620-double-F | GACAA C TTA ACG AAA CAA AAT **AAA** GAT **CGC** TAT AAA GGC TTG ACC AAT TTG G |
| Hia-618/620-R | GAC GGA GCT AGT CAG CGG ATC GAA ATT CG |
| Hia-R674A-F | GAA TAT CAC GAT CAA GTT **GCC** AAT GCG AAC GAA GTG AAA TTC |
| Hia-R674A-R | CGT TGA GCC GCC TGT GGT TTT GTC C |
| Hia-screen-F | GGC AAG AAC TTA AAA GTG AAA CAA GAG G |
| Hia-screen-R | CCA GAA CCT TTG TTG GTT ATG GTA GC |
| HiaBD1-F | AGT CAG CATATG AAC AAC AAT ACT CCT GTT ACG AAT AAG TTG |
| HiaBD1-R | AGCTAG GGATCC GCC ATT TTG ACC ATT AGC ATC GGT TG |

Point mutations are shown in **bold underlined** in the respective primers
