## Supplementary Table 3 for "The non-typeable *Haemophilus influenzae* major adhesin Hia is a dual function lectin that binds to human-specific respiratory tract sialic acid glycan receptors"

**Supplementary Table S3 -** Supplementary glycan microarray document based on MIRAGE guidelines DOI: 10.1093/glycob/cww118.

| **Classification** | **Guidelines** |
| --- | --- |
| 1. **Sample: Glycan Binding Sample** | |
| Description of Sample | Sample names:  *E. coli* BL21  *E. coli* BL21 expressing non-typeable *Haemophilus influenzae* Hia  Method of preparation:  The preparation of samples is explained in the Materials and Methods section. |
| Sample modifications | Bacteria were labelled with fluorescent dye as per methods. |
| Assay protocol | Please see *Materials and Methods*. |
| **2.** **Glycan Library** | |
| Glycan description for defined glycans | Glycans in this study are listed in Supplementary table 2 and is a published library in doi: 10.1371/journal.pntd.0004120. |
| Glycan description for undefined glycans | N/A. |
| Glycan modifications | Glycans were prepared in one of two ways for printing:  1. Glycans (with IDs in number/letter format; e.g. 1A, 4C, 7K) were sourced commercially from Dextra Laboratories, Elicityl and Carbosythn and were made into glycoamines using the protocol published in Day et al 2009 (doi: 10.1371/journal.pone.0004927).  2. Glycans (with IDs in number only format) were obtained from Prof Nicolai Bovin and were modified with spacers as per DOI: 10.1073/pnas.0407902101. The library of these glycans was first published in DOI: 10.1016/j.molimm.2009.06.010 |
| 1. **3.** **Printing Surface; e.g., Microarray Slide** | |
| Description of surface | Epoxy activated glass microarray slides. |
| Manufacturer | ArrayIt SuperEpoxy 3 (SME3). |
| Custom preparation of surface | N/A. |
| Non-covalent Immobilisation | N/A. |
| **4. Arrayer (Printer)** | |
| Description of Arrayer | SpotBot® Extreme Protein Microarray Spotter (ArrayIt, California, USA). |
| Dispensing mechanism | Contact printing using 946NS6 pins with a 6 pin in a 3 columns x 2 rows configuration. |
| Glycan deposition | Approximately 1.8 nl per spot is printed according to manufactures guidelines.  Glycan were at 500 µM in 50:50 DMF:DMSO. |
| Printing conditions | Array were printed with dehumidification at a maximum humidity of 60% relative humidity (Standard laboratory starting humidity of 75-90%) at 22ºC. Glycans were left to react with the slide for at least 8 hours after the print was completed. |
| 2. **5.** **Glycan Microarray with “Map”** | |
| Array layout | The array consists of a single array of glycans split between 6 pins (3 columns x 2 rows) with 4500μm row and column spacing. Each pin printed a 20 columns x 16 rows with 200μm spot spacing (centre to centre) with a minimum spot size of 100μm. Each sample is printed in quadruplicate with each of the 6 print areas including at least three negative control samples (print solution only) and two positive control samples consisting of one sample of fluoroscienamine and one sample of a mixture of rabbit anti-mouse antibody labeled with Alexa 555 and Alexa 647. Positive controls provide proof of successful immobilization of the amine reagents and provides for orientation for analysis. The antibodies also can provide controls for secondary antibodies used in experiments (if applicable). |
| Glycan identification and quality control | Arrays are quality controlled by a range of measures. 1. Each printed array is post print scanned to confirm deposition of the glycans on the array surface prior to neutralization of the remaining slide surface. 2. Post neutralized slides are scanned again to monitor for remaining autofluorescence. 3. Slides are assayed with fluorescently labeled lectins: WGA-Texas Red (EY Laboratories) and ConA-FITC (EY Laboratories). |
| 3. **6. Detector and Data Processing** | |
| Scanning hardware | Innoscann 9100AL. |
| Scanner settings | Scanning resolution: 10µM  Laser channel: 595nM excitation / 625nM emission filter.  PMT: 10% gain  Scan powers: Low laser power. |
| Image analysis software | ScanArray Express (Perkin Elmer). |
| Data processing | Data was exported as a CSV file and exported to Microsoft Excel. |
| **7.** **Glycan Microarray Data Presentation** | |
| Data presentation | Data is presented as yes/no binding in Figure 1. Data including glycan identification and analysed values of above background binding (see 8. below) are presented in Supplementary dataset 1. |
| 4. **8.** **Interpretation and** **Conclusion from Microarray Data** | |
| Data interpretation | We only use glycan arrays as a yes/no binding tool. Binding was determined by a 2-fold or more above the background *E.coli* BL21 control. |
| Conclusions | Hia binds to a broad range of negatively charged glycans, with the strongest binding to glycans with negative charge (Neu5Ac or sulfation) at the 6^th^ carbon of galactose. |
