## Supplementary figures and images for "The non-typeable *Haemophilus influenzae* major adhesin Hia is a dual function lectin that binds to human-specific respiratory tract sialic acid glycan receptors"

### Supplementary Figure 1

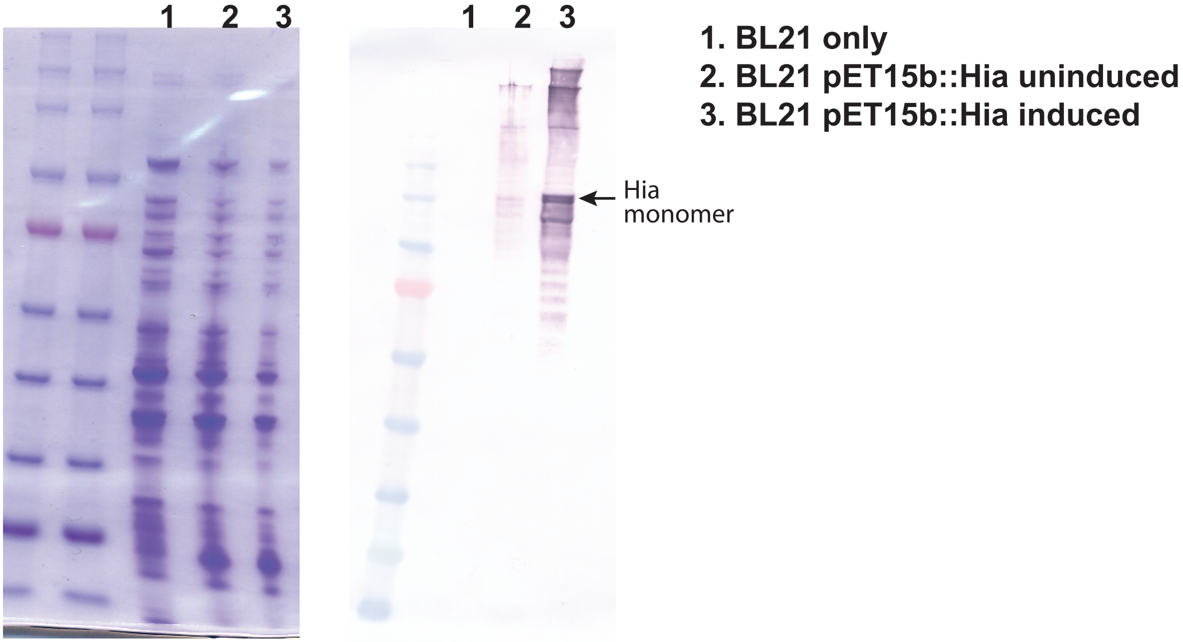

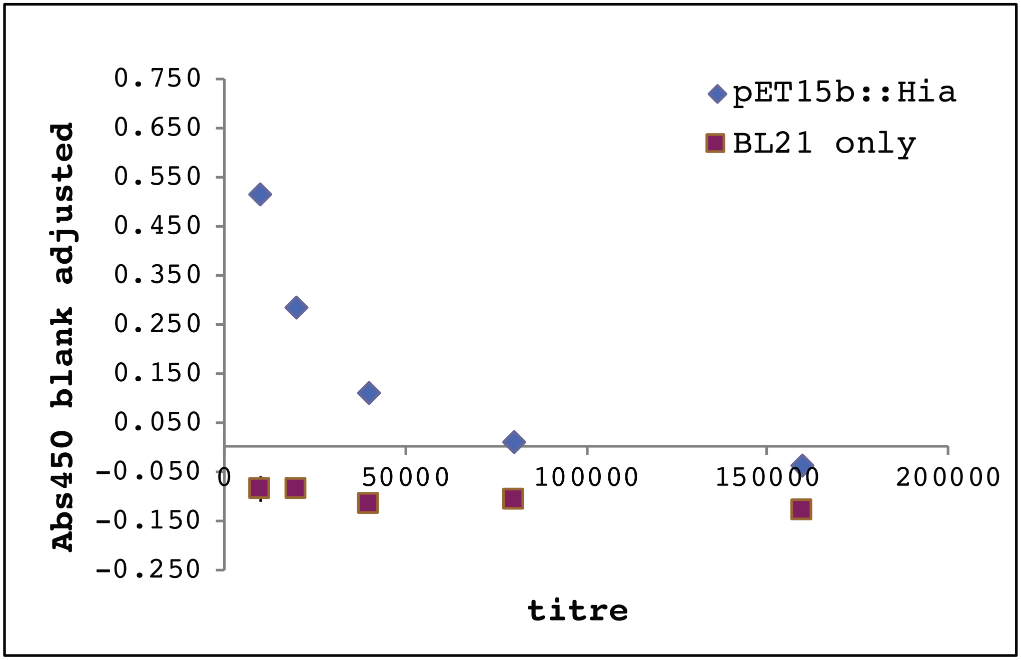
