## Supplementary Table 2 for "The non-typeable *Haemophilus influenzae* major adhesin Hia is a dual function lectin that binds to human-specific respiratory tract sialic acid glycan receptors"

| ID | Structure | Hia fold over BL21 control |
|---|---|---|
| MONOSACCHARIDES | | |
| 1 | Fucα-sp3 | 0.42 |
| 2 | Galα-sp3 | 0.43 |
| 3 | Galβ-sp3 | 0.60 |
| 4 | GalNAcα-sp0 | 0.45 |
| 5 | GalNAcα-sp3 | 0.37 |
| 6 | GalNAcβ-sp3 | 0.40 |
| 7 | Glcα-sp3 | 0.74 |
| 9 | Glcβ-sp3 | 0.26 |
| 10 | GlcNAcβ-sp3 | 0.30 |
| 14 | GlcN(Gc)β-sp4 | 0.19 |
| 15 | $HOCH_2(HOCH)_4CH_2NH_2$ | 0.47 |
| 16 | Manα-sp3 | 0.51 |
| 18 | Manβ-sp4 | 0.51 |
| 19 | ManNAcβ-sp4 | 0.53 |
| 20 | Rhaα-sp3 | 0.35 |
| 22 | GlcNAcβ-sp4 | 0.25 |
| 37 | 3-O-Su-Galβ-sp3 | 0.41 |
| 38 | 3-O-Su-GalNAcα-sp3 | 0.50 |
| 43 | 6-O-Su-GlcNAcβ-sp3 | 0.44 |
| 44 | GlcAα-sp3 | 0.33 |
| 45 | GlcAβ-sp3 | 0.36 |
| 46 | 6-$H_2PO_3$Glcβ-sp4 | 0.43 |
| 47 | 6-$H_2PO_3$Manα-sp3 | 3.92±0.61 |
| 48 | Neu5Acα-sp3 | 6.22±0.93 |
| 49 | Neu5Acα-sp9 | 5.61±0.97 |
| 52 | Neu5Gcα-sp3 | 0.34 |
| 54 | 9-NAc-Neu5Acα-sp3 | 0.54 |
| 55 | 3-O-Su-GlcNAcβ-sp3 | 0.60 |
| Terminal Galactose | | |
| 75 | Galα1-2Galβ-sp3 | 1.05 |
| 76 | Galα1-3Galβ-sp3 | 1.21 |
| 77 | Galα1-3GalNAcβ-sp3 | 1.06 |
| 78 | Galα1-3GalNAcα-sp3 | 0.97 |
| 80 | Galα1-3GlcNAcβ-sp3 | 1.20 |
| 81 | Galα1-4GlcNAcβ-sp3 | 1.08 |
| 83 | Galα1-6Glcβ-sp4 | 1.06 |

| | | |
|---|---|---|
| 84 | Galβ1-2Galβ-sp3 | 0.99 |
| 85 | Galβ1-3GlcNaAcβ-sp3 | 1.24 |
| 87 | Galβ1-3Galβ-sp3 | 1.01 |
| 88 | Galβ1-3GalNAcβ-sp3 | 1.13 |
| 89 | Galβ1-3GalNAcα-sp3 | 1.23 |
| 93 | Galβ1-4Glcβ-sp4 | 0.86 |
| 94 | Galβ1-4Galβ-sp4 | 1.07 |
| 97 | Galβ1-4GlcNAcβ-sp3 | 1.05 |
| 100 | Galβ1-6Galβ-sp4 | 1.05 |
| 145 | Galβ1-3(6-O-Su)GlcNAcβ-sp3 | 2.19±0.44 |
| 146 | Galβ1-4(6-O-Su)Glcβ-sp2 | 1.24 |
| 147 | Galβ1-4(6-O-Su)GlcNAcβ-sp3 | 2.41±0.39 |
| 150 | 3-O-Su-Galβ1-3GalNAcα-sp3 | 0.96 |
| 151 | 6-O-Su-Galβ1-3GalNAcα-sp3 | 0.90 |
| 152 | 3-O-Su-Galβ1-4Glcβ-sp2 | 1.02 |
| 153 | 6-O-Su-Galβ1-4Glcβ-sp2 | 0.94 |
| 155 | 3-O-Su-Galβ1-3GlcNAcβ-sp3 | 0.94 |
| 157 | 3-O-Su-Galβ1-4GlcNAcβ-sp3 | 0.88 |
| 159 | 4-O-Su-Galβ1-4GlcNAcβ-sp3 | 0.84 |
| 161 | 6-O-Su-Galβ1-3GlcNAcβ-sp3 | 0.98 |
| 163 | 6-O-Su-Galβ1-4GlcNAcβ-sp3 | 0.83 |
| 176 | 3-O-Su-Galβ1-4(6-O-Su)Glcβ-sp2 | 0.77 |
| 177 | 3-O-Su-Galβ1-4(6-O-Su)GlcNAcβ-sp2 | 0.97 |
| 178 | 6-O-Su-Galβ1-4(6-O-Su)Glcβ-sp2 | 1.00 |
| 179 | 6-O-Su-Galβ1-3(6-O-Su)GlcNAcβ-sp2 | 0.89 |
| 180 | 6-O-Su-Galβ1-4(6-O-Su)GlcNAcβ-sp2 | 0.63 |
| 181 | 3,4-O-$Su_2$-Galβ1-4GlcNAcβ-sp3 | 0.92 |
| 182 | 3,6-O-$Su_2$-Galβ1-4GlcNAcβ-sp2 | 0.92 |
| 183 | 4,6-O-$Su_2$-Galβ1-4GlcNAcβ-sp2 | 0.88 |
| 184 | 4,6-O-$Su_2$-Galβ1-4GlcNAcβ-sp3 | 1.18 |
| 189 | 3,6-O-$Su_2$-Galβ1-4(6-O-Su)GlcNAcβ-sp2 | 0.60 |
| 201 | 3,4-O-$Su_2$-Galβ1-4GlcNAcβ-sp3 | 0.91 |
| 203 | Galβ1-4(6-O-Su)GlcNAcβ-sp2 | 0.87 |
| 220 | Galα1-3Galβ1-4Glcβ-sp2 | 0.78 |
| 222 | Galα1-3Galβ1-4GlcNAcβ-sp3 | 0.89 |
| 224 | Galα1-4Galβ1-4Glcβ-sp3 | 0.89 |
| 225 | Galα1-4Galβ1-4GlcNAc-sp2 | 0.93 |
| 228 | Galβ1-2Galα1-4GlcNAcβ-sp4 | 0.88 |

| | | |
|---|---|---|
| 229 | Galβ1-3Galβ1-4GlcNAcβ-sp4 | 0.73 |
| 231 | Galβ1-4GlcNAcβ1-3GalNAcα-sp3 | 0.77 |
| 232 | Galβ1-4GlcNAcβ1-6GalNAcα-sp3 | 1.21 |
| 254 | Galβ1-3(GlcNAcβ1-6)GalNAcα-sp3 | 1.16 |
| 262 | Galβ1-3GalNAcβ1-3Gal-sp4 | 0.80 |
| 264 | Galβ1-4Galβ1-4GlcNAc-sp3 | 0.99 |
| 373 | Galα1-3Galβ1-4GlcNAcβ1-3Galβ-sp3 | 1.21 |
| 375 | Galα1-4GlcNAcβ1-3Galβ1-4GlcNAcβ-sp3 | 0.96 |
| 376 | Galβ1-3GlcNAcβ1-3Galβ1-4Glcβ-sp4 | 1.19 |
| 377 | Galβ1-3GlcNAcβ1-3Galβ1-3GlcNAcβ-sp2 | 1.02 |
| 378 | Galβ1-3GlcNAcα1-3Galβ1-4GlcNAcβ-sp3 | 1.20 |
| 379 | Galβ1-3GlcNAcβ1-3Galβ1-4GlcNAcβ-sp3 | 1.24 |
| 380 | Galβ1-3GlcNAcα1-6Galβ1-4GlcNAcβ-sp2 | 0.77 |
| 381 | Galβ1-3GlcNAcβ1-6Galβ1-4GlcNAcβ-sp2 | 1.04 |
| 382 | Galβ1-3GalNAcβ1-4Galβ1-4Glcβ-sp3 | 1.10 |
| 383 | Galβ1-4GlcNAcβ1-3Galβ1-4Glcβ-sp2 | 1.14 |
| 385 | Galβ1-4GlcNAcβ1-3Galβ1-4GlcNAcβ-sp3 | 1.05 |
| 387 | Galβ1-4GlcNAcβ1-6Galβ1-4GlcNAcβ-sp2 | 1.12 |
| 388 | Galβ1-4GlcNAcβ1-6(Galβ1-3)GalNAcα-sp3 | 1.14 |
| 504 | $(A\text{-}GN\text{-}M)_2$-3,6-M-GN-GNβ-sp4 | 0.88 |
| 1A | Galβ1-3Glc*N* Ac | 1.26 |
| 1B | Galβ1-4Glc*N* Ac | 1.04 |
| 1C | Galβ1-4Gal | 1.11 |
| 1D | Galβ1-6Glc*N* Ac | 1.16 |
| 1E | Galβ1-3Gal*N* Ac | 1.04 |
| 1F | Galβ1-3GalNAcβ1-4Galβ1-4Glc | 1.11 |
| 1G | Galβ1-3Glc*N* Acβ1-3Galβ1-4Glc | 0.91 |
| 1H | Galβ1-4Glc*N* Acβ1-3Galβ1-4Glc | 1.10 |
| 1I | Galβ1-4Glc*N* Acβ1-6(Galβ1-4Glc*N* Acβ1-3)Galβ1-4Glc | 1.05 |
| 1J | Galβ1-4Glc*N* Acβ1-6(Galβ1-3Glc*N* Acβ1-3)Galβ1-4Glc | 1.02 |
| 1K | Gala1-4Galβ1-4Glc | 0.99 |
| 1L | Gal*N* Acα1-*O* -Ser | 1.04 |
| 1M | Galb1-3Gal*N* Acα1-*O* -Ser | 1.02 |
| 1N | Gala1-3Gal | 0.77 |
| 1O | Gala1-3Galβ1-4Glc*N* Ac | 1.04 |
| 1P | Gala1-3Galβ1-4Glc | 0.98 |
| 2A | Gala1-3Galβ1-4Galα1-3Gal | 1.22 |
| 2B | Galβ1-6Gal | 1.20 |

| | | |
|---|---|---|
| 2C | Gal*N* Acβ1-3Gal | 1.03 |
| 2D | Gal*N* Acβ1-4Gal | 0.93 |
| 2E | Galα1-4Galβ1-4Glc*N* Ac | 1.03 |
| 2F | GalNAcα1-3Galβ1-4Glc | 1.01 |
| 2G | Galβ1-3Glc*N* Acβ1-3Galβ1-4Glc*N* Acβ1-6(Galβ1-3Glc*N* Acβ1-3)Galβ1-4Glc | 0.82 |
| 2H | Galβ1-3Glc*N* Acβ1-3Galβ1-4Glc*N* Acβ1-3Galβ1-4Glc | 0.99 |
| 18B | Galβ1-3GalNAcβ1-3Galα1-4Galβ1-4Glc | 0.85 |
| 18C | Galβ1-3GalNAcβ1-3Gal | 1.12 |
| 18L | Galβ1-4Glc | 1.23 |
| 18M | Galβ1-4Gal | 0.83 |
| 18N | Galβ1-6Gal | 1.00 |
| **Terminal N-Acetylgalactosamine** | | |
| 101 | GalNAcα1-3GalNAcβ-sp3 | 1.05 |
| 102 | GalNAcα1-3Galβ-sp3 | 0.95 |
| 103 | GalNAcα1-3GalNAcα-sp3 | 1.17 |
| 104 | GalNAcβ1-3Galβ-sp3 | 1.16 |
| 106 | GalNAcβ1-4GlcNAcβ-sp3 | 1.12 |
| 192 | GalNAcβ1-4(6-O-Su)GlcNAcβ-sp3 | 1.10 |
| 193 | 3-O-Su-GalNAcβ1-4GlcNAcβ-sp3 | 1.23 |
| 194 | 6-O-Su-GalNAcβ1-4GlcNAcβ-sp3 | 1.17 |
| 195 | 6-O-Su-GalNAcβ1-4-(3-O-Su)GlcNAcβ-sp3 | 0.81 |
| 196 | 3-O-Su-GalNAcβ1-4(3-O-Su)-GlcNAcβ-sp3 | 1.14 |
| 197 | 3,6-O-$Su_2$-GalNAcβ1-4GlcNAcβ-sp3 | 1.22 |
| 198 | 4,6-O-$Su_2$-GalNAcβ1-4GlcNAcβ-sp3 | 1.20 |
| 199 | 4,6-O-$Su_2$-GalNAcβ1-4-(3-O-Ac)GlcNAcβ-sp3 | 0.95 |
| 200 | 4-O-Su-GalNAcβ1-4GlcNAcβ-sp3 | 1.00 |
| 202 | 6-O-Su-GalNAcβ1-4(6-O-Su)GlcNAcβ-sp3 | 0.92 |
| 204 | 4-O-Su-GalNAcβ1-4GlcNAcβ-sp2 | 1.00 |
| 238 | GalNAcβ1-4Galβ1-4Glcβ-sp3 | 1.03 |
| 389 | GalNAcβ1-3Galα1-4Galβ1-4Glcβ-sp3 | 1.07 |
| 1L | Gal*N* Acα1-*O* -Ser | 1.16 |
| 2C | Gal*N* Acβ1-3Gal | 0.99 |
| 2D | Gal*N* Acβ1-4Gal | 1.10 |
| 2F | GalNAcα1-3Galβ1-4Glc | 1.21 |
| **Fucosylated** | | |
| 71 | Fucα1-2Galβ-sp3 | 1.08 |
| 72 | Fucα1-3GlcNAcβ-sp3 | 0.78 |
| 73 | Fucα1-4GlcNAcβ-sp3 | 1.10 |

| | | |
|---|---|---|
| 215 | Fucα1-2Galβ1-3GlcNAcβ-sp3 | 1.02 |
| 216 | Fucα1-2Galβ1-4GlcNAcβ-sp3 | 1.01 |
| 217 | Fucα1-2Galβ1-3GalNAcα-sp3 | 1.12 |
| 219 | Fucα1-2Galβ1-4Glcβ-sp4 | 1.07 |
| 226 | Fucα1-2(Galα1-3)Galβ-sp3 | 0.99 |
| 233 | Galβ1-3(Fucα1-4)GlcNAcβ-sp3 | 0.92 |
| 234 | Fucα1-3(Galβ1-4)GlcNAcβ-sp3 | 0.91 |
| 235 | Fucα1-2(GalNAcα1-3)Galβ-sp3 | 1.10 |
| 287 | 3-O-Su-Galβ1-3(Fucα1-4)GlcNAcβ-sp3 | 0.95 |
| 288 | Fucα1-3(3-O-Su-Galβ1-4)GlcNAcβ-sp3 | 1.03 |
| 359 | Fucα1-2(Galα1-3)Galβ1-3GlcNAcβ-sp3 | 1.09 |
| 360 | Fucα1-2(Galα1-3)Galβ1-4GlcNAcβ-sp3 | 1.19 |
| 362 | Fucα1-2(Galα1-3)Galβ1-3GalNAcα-sp3 | 0.68 |
| 363 | Fucα1-2(Galα1-3)Galβ1-3GalNAcβ-sp3 | 1.17 |
| 364 | Fucα1-3(Galα1-3Galβ1-4)GlcNAcβ-sp3 | 0.86 |
| 366 | Fucα1-2(GalNAcα1-3)Galβ1-3GlcNAcβ-sp3 | 0.98 |
| 368 | Fucα1-2(GalNAcα1-3)Galβ1-4GlcNAcβ-sp3 | 1.12 |
| 371 | Fucα1-2Galβ1-3(Fucα1-4)GlcNAcβ-sp3 | 0.82 |
| 372 | Fucα1-3(Fucα1-2Galβ1-4)GlcNAcβ-sp3 | 0.90 |
| 392 | Fucα1-2(GalNAcα1-3)Galβ1-3GalNAcα-sp3 | 1.15 |
| 479 | Fucα1-2Galβ1-3GlcNAcβ1-3Galβ1-4Glcβ-sp4 | 1.06 |
| 480 | Fucα1-2Galβ1-3GlcNAcβ1-3Galβ1-4GlcNAcβ-sp2 | 1.24 |
| 483 | Fucα1-3(Fucα1-2 (Galα1-3)Galβ1-4)GlcNAcβ-sp3 | 0.83 |
| 496 | Fucα1-2Galβ1-3(Fucα1-4)GlcNAcβ1-3Galβ1-4Glcβ-sp4 | 0.90 |
| 497 | Fucα1-3(Fucα1-2Galβ1-4)GlcNAcβ1-3Galβ1-4Glcβ-sp4 | 1.00 |
| 538 | Le$^{x}$1-6'(Le$^{c}$1-3')Lac-sp4 | 0.98 |
| 539 | LacNAc1-6'(Le$^{d}$1-3')Lac-sp4 | 0.90 |
| 541 | Le$^{x}$1-6'(Le$^{d}$1-3')Lac-sp4 | 1.08 |
| 542 | Le$^{c}$Le$^{x}$1-6'(Le$^{c}$1-3')Lac-sp4 | 0.68 |
| 543 | Le$^{x}$1-6'(Le$^{b}$1-3')Lac-sp4 | 0.86 |
| 7A | Fucα1-2Galβ1-3Glc*N* Acβ1-3Galβ1-4Glc | 1.13 |
| 7B | Galβ1-3(Fuca1-4)Glc*N* Acβ1-3Galβ1-4Glc | 0.83 |
| 7C | Galβ1-4(Fuca1-3)Glc*N* Acβ1-3Galβ1-4Glc | 0.87 |
| 7D | Fucα1-2Galβ1-3(Fucα1-4)Glc*N* Acβ1-3Galβ1-4Glc | 1.23 |
| 7E | Galβ1-3(Fucα1-4)Glc*N* Acβ1-3Galβ1-4(Fucα1-3)Glc | 1.14 |
| 7F | Fucα1-2Gal | 0.73 |
| 7G | Fucα1-2Galβ1-4Glc | 1.00 |
| 7H | Galβ1-4(Fucα1-3)Glc | 0.94 |

| | | |
|---|---|---|
| 7I | Galβ1-4(Fucα1-3)Glc*N* Ac | 0.90 |
| 7J | Galβ1-3(Fucα1-4)Glc*N* Ac | 0.99 |
| 7K | Gal*N* Acα1-3(Fucα1-2)Gal | 0.85 |
| 7L | Fucα1-2Galβ1-4(Fucα1-3)Glc | 0.81 |
| 7M | Galβ1-3(Fucα1-2)Gal | 0.91 |
| 7N | Fucα1-2Galβ1-4(Fucα1-3)Glc*N* Ac | 1.17 |
| 7O | Fucα1-2Galβ1-3Glc*N* Ac | 1.09 |
| 7P | Fucα1-2Galβ1-3(Fucα1-4)Glc*N* Ac | 1.01 |
| 8A | $SO_3$-3Galβ1-3(Fucα1-4)Glc*N* Ac | 0.99 |
| 8B | $SO_3$-3Galβ1-4(Fucα1-3)Glc*N* Ac | 0.97 |
| 8C | Galβ1-3Glc*N* Acβ1-3Galβ1-4(Fucα1-3)Glc*N* Acβ1-3Galβ1-4Glc | 0.76 |
| 8D | Galβ1-4(Fucα1-3)Glc*N* Acβ1-6(Galβ1-3Glc*N* Acβ1-3)Galβ1-4Glc | 1.01 |
| 8E | Galβ1-4(Fucα1-3)Glc*N* Acβ1-6(Fucα1-2Galβ1-3Glc*N* Acβ1-3)Galβ1-4Glc | 0.97 |
| 8F | Galβ1-4(Fucα1-3)Glc*N* Acβ1-6(Fucα1-2Galβ1-3(Fucα1-4)Glc*N* Acβ1-3)Galβ1-4Glc | 0.90 |
| 8G | Galβ1-4GlcNAcβ1-3Galβ1-4(Fucα1-3)Glc | 1.01 |
| 8H | Fucα1-2Galβ1-4(Fucα1-3)GlcNAcβ1-3Galβ1-4Glc | 0.76 |
| 8I | Fucα1-3Galβ1-4GlcNAcβ1-3Galβ1-4(Fucα1-3)Glc | 1.11 |
| 8J | Fuca1-2Galb1-4(Fuca1-3)GlcNAcb1-3(Fuca1-2)Galb1-4Glc | 0.76 |
| 8K | Galβ1-4(Fucα1-3)GlcNAcβ1-6(Galβ1-4GlcNAcβ1-3)Galβ1-4Glc | 1.17 |
| 8L | Galb1-4(Fuca1-3)GlcNAcb1-6(Galb1-4(Fuca1-3)GlcNAcb1-3)Galb1-4Glc | 0.86 |
| 8M | Fuca1-2Galb1-4(Fuca1-3)GlcNAcb1-6(Galb1-4GlcNAcb1-3)Galb1-4Glc | 0.98 |
| 8N | Galb1-3GlcNAcb1-3Galb1-4(Fuca1-3)GlcNAcb1-6(Galb1-3GlcNAcb1-3)Galb1-4Glc | 0.66 |
| 8O | Fucα1-2Galβ1-3GlcNAcβ1-3Galβb1-4(Fucα1-3)GlcNAcβ1-6(Galβ1-3GlcNAcβ1-3)Galβ1-4Glc | 0.94 |
| 8P | GalNAcb1-3(Fuca1-2)Galb1-4Glc | 1.00 |
| 9A | Galb1-3(Fuca1-2)Galb1-4(Fuca1-3)Glc | 1.22 |
| 9B | Galβ1-4Glc*N* Acβ1-6(Fucα1-2Galβ1-3Glc*N* Acβ1-3)Galβ1-4Glc | 0.81 |
| 18D | Galα1-3(Fucα1-2)Galβ1-4Glc | 0.96 |
| 18E | GalNAcα1-3(Fucα1-2)Galβ1-4(Fucα1-3)Glc | 0.87 |
| 19J | Galβ1-4(Fucα1-3)GlcNAcβ1-3Gal | 1.02 |
| 19L | Fucα1-2Galβ1-4(Fucα1-3)GlcNAcβ1-3Gal | 0.93 |
| 19M | Galβ1-3(Fucα1-4)GlcNAcβ1-3Gal | 0.87 |
| 19N | Fucα1-2Galβ1-3(Fucα1-4)GlcNAcβ1-3Gal | 0.70 |
| **Sialylated** | | |
| 169 | Neu5Acα2-3Galβ-sp3 | 1.29 |
| 170 | Neu5Acα2-6Galβ-sp3 | 8.84±1.47 |
| 171 | Neu5Acα2-3GalNAcα-sp3 | 1.30 |
| 172 | Neu5Acα2-6GalNAcα-sp3 | 1.95 |
| 174 | Neu5Gcα2-6GalNAcα-sp3 | 1.58 |

| | | |
|---|---|---|
| 186 | Neu5Acα2-8Neu5Acα2-sp3 | 1.78 |
| 205 | Neu5Acα2-6GalNAcβ-sp3 | 9.34±1.69 |
| 206 | Neu5Gcα2-3Gal-sp3 | 1.27 |
| 289 | Galα1-3(Neu5Acα2-6)GalNAcα-sp3 | 1.36 |
| 290 | Galβ1-3(Neu5Acα2-6) GalNAcα-sp3 | 1.19 |
| 292 | Neu5Acα2-3Galβ1-3GalNAcα-sp3 | 1.60 |
| 293 | Neu5Acα2-3Galβ1-4Glcβ-sp3 | 1.59 |
| 294 | Neu5Acα2-3Galβ1-4Glcβ-sp4 | 1.69 |
| 295 | Neu5Acα2-6Galβ1-4Glcβ-sp2 | 1.56 |
| 298 | Neu5Acα2-3Galβ1-4GlcNAcβ-sp3 | 3.34±0.89 |
| 299 | Neu5Acα2-3Galβ1-3GlcNAcβ-sp3 | 1.81 |
| 300 | Neu5Acα2-6Galβ1-4GlcNAcβ-sp3 | 1.82 |
| 303 | Neu5Gcα2-3Galβ1-4GlcNAcβ-sp3 | 1.33 |
| 304 | Neu5Gcα2-6Galβ1-4GlcNAcβ-sp3 | 1.45 |
| 306 | 9-NAc-Neu5Acα2-6Galβ1-4GlcNAcβ-sp3 | 1.29 |
| 315 | Neu5Acα2-3Galβ1-4-(6-O-Su)GlcNAcβ-sp3 | 1.84 |
| 317 | Neu5Acα2-3Galβ1-3-(6-O-Su)GalNAcβ-sp3 | 1.57 |
| 318 | Neu5Acα2-6Galβ1-4-(6-O-Su)GlcNAcβ-sp3 | 9.44±2.21 |
| 319 | Neu5Acα2-3-(6-O-Su)Galβ1-4GlcNAcβ-sp3 | 4.37±0.67 |
| 321 | (Neu5Acα2-8)$_3$-sp3 | 1.38 |
| 323 | Neu5Acα2-6Galβ1-3GlcNAc-sp3 | 1.20 |
| 324 | Neu5Acα2-6Galβ1-3(6-O-Su)GlcNAc-sp3 | 1.32 |
| 331 | Neu5Gcα2-3Galβ1-3GlcNAcβ-sp3 | 1.48 |
| 421 | Neu5Acα2-3(GalNAcβ1-4)Galβ1-4Glcβ-sp2 | 1.26 |
| 422 | Neu5Acα2-3Galβ1-4GlcNAcβ1-3Galβ-sp3 | 1.33 |
| 423 | Fucα1-3(Neu5Acα2-3Galβ1-4)GlcNAcβ-sp3 | |
| 426 | Neu5Acα2-3Galβ1-3(Fucα1-4)GlcNAcβ-sp3 | 1.28 |
| 428 | Fucα1-3(Neu5Acα2-3Galβ1-4)6-O-Su-GlcNAcβ-sp3 | 1.40 |
| 429 | Fucα1-3(Neu5Acα2-3(6-O-Su)Galβ1-4)GlcNAcβ-sp3 | 1.39 |
| 433 | Neu5Acα2-3Galβ1-3(Neu5Acα2-6)GalNAcα-sp3 | 1.31 |
| 434 | Neu5Acα2-8Neu5Acα2-3Galβ1-4Glcβ-sp4 | 1.22 |
| 527 | Neu5Acα2-3Galβ1-4GlcNAcβ1-3Galβ1-4GlcNAcβ-sp2 | 1.34 |
| 528 | Fucα1-3(Neu5Acα2-3 Galβ1-4)GlcNAcβ1-3Galβ-sp3 | 4.33±1.12 |
| 529 | Neu5Acα2-6(Galβ1-3)GlcNAcβ1-3Galβ1-4Glcβ-sp4 | 1.46 |
| 531 | GalNAcβ1-4(Neu5Acα2-8Neu5Acα2-3)Galβ1-4Glc-sp2 | 1.29 |
| 532 | Neu5Acα2-8Neu5Acα2-8Neu5Acα2-3Galβ1-4Glc-sp2 | 1.48 |
| 533 | (Neu5Acα2-8)2Neu5Acα2-3(GalNAcβ1-4)Galβ1-4Glc-sp2 | 1.55 |
| 534 | Neu5Acα2-3Galβ1-4GlcNAcβ1-3Galβ1-4GlcNAcβ-sp3 | 1.47 |

| | | |
|---|---|---|
| 536 | Neu5Acα2-3Galβ1-3GlcNAcβ1-3Galβ1-4Glcβ-sp4 | 1.86 |
| 537 | Neu5Acα2-3Galβ1-4GlcNAcβ1-3Galβ1-4Glcβ-sp4 | 1.77 |
| 540 | $Le^x$1-6'(6'SLN1-3')Lac-sp4 | 1.56 |
| 10A | Neu5Acα2-3Galβ1-3(Fucα1-4)Glc*N* Ac | 1.62 |
| 10B | Neu5Acα2-3Galβ1-4(Fucα1-3)Glc*N* Ac | 1.62 |
| 10C | Neu5Acα2-3Galβ1-3Glc*N* Acβ1-3Galβ1-4Glc | 1.66 |
| 10D | Galβ1-4(Fucα1-3)Glc*N* Acβ1-6(Neu5Acα2-6Galβ1-4Glc*N* Acβ1-3)Galβ1-4Glc | 1.45 |
| 10E | Neu5Acα2-3Galβ1-3(Neu5Acα2-6)GalNAc | 1.33 |
| 10H | Neu5Acα2-6Galβ1-3GlcNAcβ1-3Galβ1-4(Fucα1-3)Glc | 7.47±1.42 |
| 10I | Galβ1-3GlcNAcβ1-3(Neu5Acα2-6Galβ1-4GlcNAcβ1-6)Galβ1-4Glc | 1.51 |
| 10J | Neu5Acα2-6Galβ1-3GlcNAcβ1-3(Galβ1-4GlcNAcβ1-6)Galβ1-4Glc | 1.59 |
| 10K | Neu5Acα2-3Galβ1-4Glc*N* Ac | 1.18 |
| 10L | Neu5Acα2-6Galβ1-4Glc*N* Ac | 11.56±2.37 |
| 10M | Neu5Acα2-3Galβ1-3GlcNAcβ1-3Galβ1-4Glc | 1.49 |
| 10N | Galβ1-3(Neu5Acα2-6)Glc*N* Acβ1-3Galβ1-4Glc | 1.32 |
| 10O | Neu5Acα2-6Galβ1-4Glc*N* Acβ1-3Galβ1-4Glc | 1.97 |
| 10P | Neu5Acα2-3Galβ1-3(Neu5Acα2-6)Glc*N* Acβ1-3Galβ1-4Glc | 1.80 |
| 11A | Neu5Acα2-3Galβ1-4Glc | 1.67 |
| 11B | Neu5Acα2-6Galβ1-4Glc | 1.94 |
| 11C | (Neu5Acα2-8Neu5Ac)n (n<50) | 1.29 |
| 11D | Neu5Acα2-6Galβ1-4GlcNAcβ1-2Manα1-6(Neu5Acα2-6Galβ1-4GlcNAcβ1-2Manα1-6)Manβ1-4GlcNAcβ1-4GlcNAc-Asn | 1.90 |
| 18A | Neu5Acα2-3Galβ1-4GlcNAcβ1-3Galβ1-4Glc | 1.93 |
| 18K | 9-NAc-Neu5Ac | 1.63 |
| 18O | Neu5Gc | 1.32 |
| 19K | Neu5Acα2-3Galβ1-4(Fucα1-3)GlcNAcβ1-3Gal | 1.24 |
| | **Mannose** | |
| 119 | Manα1-2Manβ-sp4 | 0.79 |
| 120 | Manα1-3Manβ-sp4 | 1.17 |
| 121 | Manα1-4Manβ-sp4 | 0.94 |
| 122 | Manα1-6Manβ-sp4 | 1.24 |
| 123 | Manβ1-4GlcNAcβ-sp4 | 1.48 |
| 124 | Manα1-2Manα-sp4 | 0.73 |
| 258 | Manα1-3(Manα1-6)Manβ-sp4 | 1.38 |
| 495 | Manα1-3(Manα1-3(Manα1-6)Manα1-6)Manβ-sp4 | 0.92 |
| 5A | Glc*N* Acβ1-2Man | 1.36 |
| 5B | Glc*N* Acβ1-2Manα1-6(Glc*N* Acβ1-2Manα1-3)Man | 1.39 |
| 5C | Manα1-2Man | 0.92 |
| 5D | Manα1-3Man | 1.03 |

| | | |
|---|---|---|
| 5E | Manα1-4Man | 0.95 |
| 5F | Manα1-6Man | 1.48 |
| 5G | Manα1-6(Manα1-3)Man | 0.98 |
| 5H | Manα1-6(Manα1-3)Manα1-6(Manα1-3)Man | 0.93 |
| **Terminal N-Acetylglucosamine** | | |
| 113 | GlcNAcβ1-3GalNAcα-sp3 | 0.84 |
| 114 | GlcNAcβ1-3Manβ-sp4 | 1.04 |
| 115 | GlcNAcβ1-4GlcNAcβ-Asn | 0.84 |
| 117 | GlcNAcβ1-4GlcNAcβ-sp4 | 0.94 |
| 118 | GlcNAcβ1-6GalNAcα-sp3 | 0.83 |
| 149 | GlcNAcβ1-4(6-O-Su)GlcNAcβ-sp2 | 0.85 |
| 167 | GlcNAcβ1-4-[HOOC($CH_3$)CH]-3-O-GlcNAcβ-sp4 | 1.05 |
| 168 | GlcNAcβ1--[HOOC($CH_3$)CH]-3-O-GlcNAcβ-L-alanyl-D-i-glutaminyl-L-lysine | 0.92 |
| 246 | GlcNAcβ1-2Galβ1-3GalNAcα-sp3 | 0.87 |
| 247 | GlcNAcβ1-3Galβ1-3GalNAcα-sp3 | 0.82 |
| 248 | GlcNAcβ1-3Galβ1-4Glcβ-sp2 | 0.93 |
| 250 | GlcNAcβ1-3Galβ1-4GlcNAcβ-sp3 | 0.96 |
| 251 | GlcNAcβ1-4Galβ1-4GlcNAcβ-sp2 | 1.08 |
| 252 | GlcNAcβ1-4GlcNAcβ1-4GlcNAcβ-sp4 | 0.81 |
| 253 | GlcNAcβ1-6Galβ1-4GlcNAcβ-sp2 | 0.70 |
| 255 | GlcNAcβ1-3(GlcNAcβ1-6)GalNAcα-sp3 | 1.03 |
| 395 | GlcNAcβ1-3(GlcNAcβ1-6)Galβ1-4GlcNAcβ-sp3 | 1.14 |
| 493 | (GlcNAcβ1-4)$_5$β-sp4 | 1.04 |
| 503 | (GlcNAcβ1-4)$_6$β-sp4 | 1.09 |
| 505 | (GN-M)$_2$-3,6-M-GN-GNβ-sp4 | 1.21 |
| 4A | Glc*N* Acβ1-4Glc*N* Ac | 1.06 |
| 4B | Glc*N* Acβ1-4Glc*N* Acβ1-4Glc*N* Ac | 0.93 |
| 4C | Glc*N* Acβ1-4Glc*N* Acβ1-4Glc*N* Acβ1-4Glc*N* Ac | 0.74 |
| 4D | Glc*N* Acβ1-4Glc*N* Acβ1-4Glc*N* Acβ1-4Glc*N* Acβ1-4Glc*N* Acβ1-4Glc*N* Ac | 0.95 |
| 4E | Bacterial cell wall muramyl discaccharide | 0.89 |
| 4F | Glc*N* Acβ1-4Glc*N* Acβ1-4Glc*N* Acβ1-4Glc*N* Acβ1-4Glc*N* Ac | 0.97 |
| 18G | 6-O-Su-GlcNAc | 0.73 |
| 18H | GlcNAc | 1.06 |
| **Glucose** | | |
| 110 | Glcα1-4Glcβ-sp3 | 1.24 |
| 111 | Glcβ1-4Glcβ-sp4 | 1.03 |
| 112 | Glcβ1-6Glcβ-sp4 | 0.96 |
| 164 | GlcAβ1-3GlcNAcβ-sp3 | 0.93 |

| | | |
|---|---|---|
| 165 | GlcAβ1-3Galβ-sp3 | 0.85 |
| 166 | GlcAβ1-6Galβ-sp3 | 0.85 |
| 240 | (Glcα1-4)$_3$β-sp4 | 0.97 |
| 241 | (Glcα1-6)$_3$β-sp4 | 0.90 |
| 390 | (Glcα1-4)$_4$β-sp4 | 0.76 |
| 391 | (Glcα1-6)$_4$β-sp4 | 0.87 |
| 492 | (Glcα1-6)$_5$β-sp4 | 0.87 |
| 502 | (Glcα1-6)$_6$β-sp4 | 0.66 |
| 18I | GlcA | 0.84 |
| 18J | 6-O-($H_2PO_4$)-Glc | 1.08 |
| 19O | Glcα1-4Glcα1-4 | 0.84 |
| 19P | Glcα1-4Glcα1-4Glcα1-4 | 0.84 |
| **Low molecular weight Carageenan and Glycoaminoglycans (GAGS)** | | |
| 12A | Neocarratetraose-41, 3-di-O-sulphate (Na+) | 0.93 |
| 12B | Neocarratetraose-41-O-sulphate (Na+) | 2.36±0.67 |
| 12C | Neocarrahexaose-24,41, 3, 5-tetra-O-sulphate (Na+) | 1.08 |
| 12D | Neocarrahexaose-41, 3, 5-tri-O-sulphate (Na+) | 2.43±0.71 |
| 12E | Neocarraoctaose-41, 3, 5, 7-tetra-O-sulphate (Na+) | 0.73 |
| 12F | Neocarradecaose-41, 3, 5, 7, 9-penta-O-sulphate (Na+) | 1.03 |
| 12G | ΔUA-2S-GlcNS-6S | 1.05 |
| 12H | ΔUA-GlucNS-6S | 0.96 |
| 12I | ΔUA-2S-GlucNS | 0.86 |
| 12J | ΔUA-2S-GlcNAc-6S | 0.93 |
| 12K | ΔUA-GlcNAc-6S | 2.13±0.56 |
| 12L | ΔUA-2S-GlcNAc | 0.95 |
| 12M | ΔUA-GlcNAc | 1.16 |
| 12N | ΔUA-GalNAc-4S (Delta Di-4S) | 1.27 |
| 12O | ΔUA-GalNAc-6S (Delta Di-6S) | 0.78 |
| 12P | ΔUA-GalNAc-4S,6S (Delta Di-disE) | 0.68 |
| 13A | ΔUA-2S-GalNAc-4S (Delta Di-disB) | 0.90 |
| 13B | ΔUA-2S-GalNAc-6S (Delta Di-disD) | 0.86 |
| 13C | ΔUA-2S-GalNAc-4S-6S (Delta Di-tisS) | 0.73 |
| 13D | ΔUA-2S-GalNAc-6S (Delta Di-UA2S) | 0.87 |
| 13E | ΔUA-GlcNAc (Delta Di-HA) | 0.81 |
| 14M | ΔUA→2S-GlcN-6S | 4.32±0.86 |
| 14N | ΔUA→GlcN-6S | 0.89 |
| 14O | ΔUA→2S-GlcN | 0.95 |
| 14P | ΔUA→GlcN | 0.92 |

| | | |
|---|---|---|
| **High molecular weight Carageenan and Glycoaminoglycans (GAGS)** | | |
| 625 | (GlcAβ1-4GlcNAcβ1-3)$_8$-NH$_2$-ol | 1.02 |
| 13F | (GlcAβ1-3Glc*N* Acβ1-4)n (n=4) | 1.02 |
| 13G | (GlcAβ1-3Glc*N* Acβ1-4)n (n=8) | 1.11 |
| 13H | (GlcAβ1-3Glc*N* Acβ1-4)n (n=10) | 1.07 |
| 13I | (GlcAβ1-3Glc*N* Acβ1-4)n (n=12) | 2.49±0.52 |
| 13J | (GlcA/IdoAα/β1-4Glc*N* Acα1-4)n (n=200) | 0.88 |
| 13K | (GlcA/IdoAβ1-3(±4/6S)Gal*N* Acβ1-4)n (n<250) | 3.78±0.88 |
| 13L | ((±2S)GlcA/IdoAα/b1-3(±4S)Gal*N* Acβ1-4)n (n<250) | 0.78 |
| 13M | (GlcA/IdoAβ1-3(±6S)Gal*N* Acβ1-4)n (n<250) | 1.11 |
| 13N | HA - 4 | 2.21±0.91 |
| 130 | HA - 6 | 1.40 |
| 13P | HA - 8 | 0.97 |
| 14A | HA 10 | 0.30 |
| 14B | HA-12 | 0.83 |
| 14C | HA-14 | 1.37 |
| 14D | HA-16 | 1.43 |
| 14E | HA 30000 da | 1.35 |
| 14F | HA 107000 da | 0.78 |
| 14G | HA 190000 da | 0.60 |
| 14H | HA 220000 da | 0.71 |
| 14I | HA 1600000 da | 0.86 |
| 14J | Heparin sulfate | 4.56±1.28 |
| 14K | β1-3Glucan | 1.01 |
| **Complex N-glycans** | | |
| 627 | (Sia2-6A-GN-M)$_2$-3,6-M-GN-GNβ-sp4 | 1.69 |
| 19A | Galβ1-4GlcNAcβ1-2Manα1-3(Galβ1-4GlcNAcβ1-2Manα1-6Man)β1-4GlcNAcβ1-4(Fucα1-6)GlcNAc | 1.19 |
| 19B | Galβ1-4GlcNAcβ1-2(Galβ1-4GlcNAcβ1-4)Manα1-3(Galβ1-4GlcNAcβ1-2(Galβ1-4GlcNAcβ1-6)Manα1-6Man)β1-4GlcNAcβ1-4GlcNAc | 0.97 |
| 19C | Neu5Acα2-6Galβ1-4GlcNAcβ1-2Manα1-3(Galβ1-4GlcNAcβ1-2Manα1-6)Manβ1-4GlcNAcβ1-4GlcNAc | 1.52 |
| 19D | Neu5Acα2-6Galβ1-4GlcNAcβ1-2Manα1-3(Neu5Acα2-6Galβ1-4GlcNAcβ1-2Manα1-6)Manβ1-4GlcNAcβ1-4GlcNAc | 1.58 |
| 19E | Galβ1-4GlcNAcβ1-2Manα1-3(Galβ1-4GlcNAcβ1-2Manα1-6)Manβ1-4GlcNAcβ1-4GlcNAc | 0.84 |
| 19F | Neu5Acα2-6Galβ1-4GlcNAcβ1-2Manα1-3(Neu5Acα2-6Galβ1-4GlcNAcβ1-2Manα1-6)Manβ1-4GlcNAcβ1-4(Fucα1-6)GlcNAc | 1.82 |
| 19G | Neu5Acα2-6Galβ1-4GlcNAcβ1-2(Neu5Acα2-6Galβ1-4GlcNAcβ1-4)Manα1-3(Neu5Acα2-6Galβ1-4GlcNAcβ1-2Manα1-6)Manβ1-4GlcNAcβ1-4Glc | 1.93 |
| 19H | GlcNAcβ1-2(GlcNAcβ1-4)Manα1-3(GlcNAcβ1-2Manα1-6)GlcNAcβ1-4Manβ1-4GlcNAcβ1-4GlcNAc | 0.61 |

Red/blue indicates binding. Binding is determined by positive interaction in three replicate array experiments. Positive interactions are determined by a background subtracted fluorescence value significantly above (greater than 2-fold) BL21 not expressing Hia protein.
